# Characterization of proteome-size scaling by integrative omics reveals mechanisms of proliferation control in cancer

**DOI:** 10.1101/2022.06.21.496989

**Authors:** Ian Jones, Lucas Dent, Tomoaki Higo, Theo Roumeliotis, Mar Arias Garcia, Hansa Shree, Jyoti Choudhary, Malin Pedersen, Chris Bakal

## Abstract

Almost all living cells maintain size uniformity through successive divisions. Proteins that sub- or super-scale with size act as rheostats which regulate cell progression. A comprehensive atlas of these proteins is lacking; particularly in cancer cells where both mitogen and growth signalling are dysregulated.

Utilising a multi-omic strategy, that integrates quantitative single cell imaging, phosphoproteomic and transcriptomic datasets, we leverage the inherent size heterogeneity of melanoma cells to investigate how peptides, post-translational modifications, and mRNAs scale with cell size to regulate proliferation. We find melanoma cells have different mean sizes, but all retain uniformity. Across the proteome, we identify proteins and phosphorylation events that ‘sub’ and ‘super’ scale with cell size. In particular, G2/M, biosynthetic, and cytoskeletal regulators sub- and super-scale with size. In small cells growth and proliferation processes are tightly coupled by translation which promotes CCND1 accumulation and anabolic increases in mass. Counter intuitively, anabolic growth pathways and translational process are low in large cells, which throttles the expression of factors such as CCND1 and thereby coupling proliferation from anabolic growth. Strikingly, these cells exhibit increased growth and comparable proliferation rates. Mathematical modelling suggests that decoupling growth and proliferative signalling fosters proliferation under mitogenic inhibition. As factors which promote adhesion and actin reorganization super-scale with size or are enriched in large cells, we suggest that growth/proliferation in these cells may be decoupled by cell spreading and mechanics. This study provides one of the first demonstrations of size-scaling phenomena in cancer and how morphology determines the chemistry of the cell.

## Introduction

Eukaryotic cells vary widely in size; there is a billion-fold difference in cell volume between *Xenopus* oocytes (∼1mm diameter (Wallace et al., 1981)) and phytoplankton (∼1um) (Palenik et al., 2007). This results in a spectrum of biology, as cell size directly impacts nutrient acquisition and consumption, diffusive processes, and intracellular protein concentrations (Amodeo & Skotheim, 2016; Bernal-Mizrachi et al., 2001; Monds et al., 2014; Ruvinsky et al., 2005).

Although striking differences in size are observed when comparing between different cell types, size distributions within proliferating cell types show only modest variance in cell size, or size ‘uniformity’ (Ginzberg et al., 2015) (coefficients of variation, CVs, typically 0.1-0.3 (Scotchman et al., 2021)). The size homogeneity of proliferating cell populations implies the existence of size checkpoints that occur during proliferation and coordinate cell cycle progression and acquisition of cell mass (Amodeo & Skotheim, 2016; Ginzberg et al., 2015).

To maintain a stable size distribution across a population, a checkpoint system can measure the size of individual cells with molecular ‘rulers’ (Schmoller et al., 2015). Measurements are then coupled to the speed of the division cycle and the acquisition of mass. Such a system would ‘penalise’ cells that deviate from the target volume (a ‘sizer’ system), accelerating or diminishing the cell proliferation rate. Notably growth is not necessarily reliant on protein synthesis and anabolism (Miettinen et al., 2021).

Other mechanisms of size determination have been documented that do not inherently depend on cell size measurements, such as the ‘adder’ or ‘timer’ models where a fixed amount of cell mass is added per cycle (Amodeo & Skotheim, 2016; Campos et al., 2014). However, recent studies allude to similarity between sizer and adder/timer systems, with modest errors in sizer function leading to adder-like behaviour (Facchetti, Knapp, Chang, et al., 2019).

Several studies have identified molecular mechanisms of how size measurements are coupled to proliferation and/or growth. In budding yeast a type of ‘ruler’ appears to consist of a mechanism where the concentration of a cell cycle progression inhibitor, Whi5, becomes diluted with respect to the activator, Cln3, as cells grow larger, allowing cell cycle progression only at a critical size. (Costanzo et al., 2004; Schmoller et al., 2015). The set-point appears to be in part determined by the concentration of Whi5 is relative to the number of DNA binding sites for the cell cycle activator SBF. (Heldt et al., 2018).

Recently, RB1 (an ortholog of Whi5) has been demonstrated to have a role in mammalian size control. RB1 concentration sub-scales with size across the cell cycle. Meaning, in smaller newly born daughters, the activity of RB1 exceeds that of its agonist Cyclin D1 (CCND1). CCND1 scales with size, and thus as cells grow, there is a point at which the activity of CCND1 exceeds that over RB1 and proliferation occurs (Zatulovskiy et al., 2020). In normal cells CCDN1 levels themselves are a function of mitogen signalling and translational activity (Min et al., 2020). Thus, in normal mammalian cells, cells meet the RB1:CCDN1 set-point for proliferation by increasing the concentration of CCND1 with regards to RB1 by actively translating CCND1 while simultaneously diluting RB1 as cells grow larger (Zatulovskiy et al., 2020).

It is becoming clear that regulation of protein function by diluting or concentrating with cell size is not a rare phenomenon. Many proteins have been shown to ‘super’ or ‘sub’ scale (mass fraction increases/decreases) with cell size (Amodeo et al., 2015; Lanz et al., 2021) beyond a small set of proliferative regulators. Indeed, recent studies point to histones (Amodeo et al., 2015; Swaffer et al., 2021), translational components (Yahya et al., 2021) and several metabolic elements (Lanz et al., 2021; Neurohr et al., 2019) sub/super scaling with cell size. Not all these proteins will act as size ‘rulers’, and may instead influence their activity. For example, chromatin-associated histones have been shown to regulate equal partitioning of Whi5 in asymmetric cell divisions in budding yeast (Swaffer et al., 2021). Dilution of cell proteins (and intracellular DNA) through excessive growth has also recently been associated with the onset of cell senescence (Neurohr et al., 2019).

In other cells, size control is highly influenced by cell geometry. For example, in fission yeast, (Fantes & Nurse, 1977), size is thought to be determined primarily at G2/M through the accumulation of localized CDR2 nodes that accumulate at the growing mid body to activate CDK1 by inhibiting Wee1 (Facchetti, Knapp, Flor-Parra, et al., 2019; Lundgren et al., 1991; Russell & Nurse, 1987). Because CDR2 accumulation scales with surface area, this provides a means by which the detection of cell geometry influences a size checkpoint. Other work has shown analogous regulation of size by surface area or volume in both bacterial (Harris & Theriot, 2016), and mammalian cells (Varsano et al., 2017). Thus, different geometric quantities, such as surface area, may serve to mediate size control in different cell types through coupling to signal proteins.

Because most studies on size control use either yeast, bacteria, or normal mammalian cells, there is little understanding of size determination in cancer. Classic studies suggest that increased size and morphological heterogeneity are histological measures of cancer grade, with large and more morphologically varied cancers, tending to be more pathogenic (Watson, 1997). Indeed, only highly heterogeneously sized lines induced tumours upon transplantation in mice (Caspersson et al., 1963). This diversification of cell size has been shown to be cell autonomous and not an artefact of the environment (Caspersson et al., 1963). Together these observations suggest modification of cell size in malignant tissues and that this contributes to (or coincides with) increased cellular fitness. But the exact relationship between size and disease is poorly understood.

Consistent with the idea that size and size heterogeneity are associated with oncogenesis, dysregulation of RB1’s inhibitory actions on E2F1, a putative size ruler, are frequent oncogenic events (Nevins, 2001). For example, many cancers have loss of function mutations in the RB1 gene and/or have upregulated activity in ERK kinases, which promotes increased CCND1 levels, and a concomitant increased activation of RB1’s inhibitor CDK4/6 (Chinnam & Goodrich, 2011). Indeed, mutations resulting in constitutively active BRAF or NRAS, proteins in-part responsible for the activation of ERK kinases and ultimately CCND1 production (Joseph et al., 2010), comprise 50% and 20% of all melanoma cases, respectively (Davies et al., 2002; Reifenberger et al., 2004). These common driver mutations are likely to directly affect the size control machinery, however, the specific effect of these mutations on size control is essentially unknown.

Here we leverage the natural phenotypic heterogeneity of a panel of melanoma cell lines, to investigate the size-scaling of intracellular peptides and transcripts in the context of cell growth and division. We show that BRAF and NRAS mutant melanomas have diverse mean sizes, but size uniformity is maintained. Both RB1 and Cyclin D1 sub-scale with size across lines. However, the relative ratio of these proteins is constant, suggesting a common set-point of RB1 to Cyclin D1 is a conserved despite the presence of oncogenic mutations which can affect the levels of both proteins. We identify sub and super-scaling across the cell proteome and phosphoproteome. In particular, we show that regulators of G2/M, translation, and growth sub-scale with size across lines, but stress response proteins, adhesion components, and certain growth factor receptors super-scale. Through integration of transcriptomic data, we show that scaling of translation is regulated transcriptionally. mTOR signalling and translation couples cell growth and proliferation by promoting anabolic growth and increasing CCND1 levels in small populations. In contrast, larger lines counter intuitively have decreased levels of translation and altered biosynthetic signalling despite exhibiting an increased growth rate. These cells may spread to concentrate anabolic regulators subverting the constraints of a reduced biosynthetic mass fraction. Proteomic data suggests this could be due to reorganization of adhesion and actin structures. Mathematical modelling indicates that uncoupling growth and proliferative systems facilitates division following a reduction mitogenic signalling. This research provides one of the first datasets describing how the transcriptional and proteomic profile of melanoma cells can change with cell size, indicating that cell morphology can have direct and meaningful effects on the chemistry of the cell.

## Results

### Melanoma cell lines exhibit comparable size control but different cell sizes

To understand the relationship between cell size and different clinically relevant oncogenic drivers, we quantified the morphology of 17,547 single cells from 11 mouse melanoma cell lines from three different genetic backgrounds (**SD1, SF1**). Lines were either: BRAF*; constitutively active BRAF typically due to a V600E mutation (Dhomen et al., 2009) (Cantwell-Dorris et al., 2011); NRAS*, constitutively active NRAS, due to G12D mutations (Pedersen et al., 2013, 2014) (Burd et al., 2014) or NRAS*/KDBRAF, where lines harboured a constitutively active NRAS mutation and a dominant negative mutation in the BRAF kinase domain BRAF (D594A) (**Table 1**) (Pedersen et al., 2014). NRAS*/KDBRAF mutants mimic the clinical situation where there is paradoxical activation of BRAF following treatment of NRAS mutant cells with BRAF inhibitors such as Vemurafenib (Poulikakos et al., 2010) (Bhargava et al., 2016).

**Table 1:** Cell Line Details.

| Cell ID | Species | Genotype Details | Derived from | Publication |
| --- | --- | --- | --- | --- |
| 19161 | murine | NRAS mut/Tyrosinase CreA | Brain melanoma | Pedersen et al. Cancer Discovery 2013 |
| 19398 | murine | NRAS mut/Tyrosinase CreA | Brain melanoma | Pedersen et al. Cancer Discovery 2013 |
| C873 | murine | NRAS mut/Tyrosinase CreA | Brain melanoma | Pedersen et al. Cancer Discovery 2013 |
| Ear tum | murine | NRAS mut/BRAF KD under Tyrosinase CreERT with TAM | Cutaneous melanoma | Pedersen et al. PCMR 2014 |
| EAR B | murine | NRAS mut/BRAF KD under Tyrosinase CreERT with TAM | Cutaneous melanoma | Pedersen et al. PCMR 2014 |
| 22532 | murine | NRAS mut/Tyrosinase CreA | Brain melanoma | Pedersen et al. Cancer Discovery 2013 |
| 14508 LN (A) | murine | NRAS mut/BRAF KD under Tyrosinase CreERT with TAM | Cutaneous melanoma | Pedersen et al. PCMR 2014 |
| 17569 | murine | NRAS mut/Tyrosinase CreA | Brain melanoma | Pedersen et al. Cancer Discovery 2013 |
| 22783 | murine | NRAS mut/BRAF KD under Tyrosinase CreERT with TAM | Cutaneous melanoma | Pedersen et al. PCMR 2014 |
| 4434 (KB) | murine | BRAF mut/p16-/- | Cutaneous melanoma | Dhomen et al. Cancer Cell 2009 |
| 17864 | murine | NRAS mut/UV | Cutaneous melanoma | Pedersen unpublished |
| 17864A | murine | NRAS mut/UV | Cutaneous melanoma | Pedersen unpublished |
| 21917 | murine | NRAS mut/UV | Cutaneous melanoma | Pedersen unpublished |
| 21015 | murine | BRAF mut/PTEN null | Cutaneous melanoma | Pedersen unpublished |
| 24038 | murine | NRAS mut/BRAF KD | Cutaneous melanoma | Pedersen et al. PCMR 2014 |
| B14341 | murine | NRAS mut/BRAF KD | Cutaneous melanoma | Pedersen et al. PCMR 2014 |
| C876 | murine | NRAS mut/Tyrosinase CreA | Brain melanoma | Pedersen et al. Cancer Discovery 2013 |
| C790 | murine | NRAS mut/Tyrosinase CreA | Brain melanoma | Pedersen et al. Cancer Discovery 2013 |
| 17568 | murine | NRAS mut/Tyrosinase CreA | Brain melanoma | Pedersen et al. Cancer Discovery 2013 |
| 5555 | murine | BRAF mut/p16-/- | Cutaneous melanoma | Dhomen et al. Cancer Cell 2009 |
| NRASQ61 | murine | NRAS Q61A | Cutaneous melanoma | Pedersen unpublished |

For each single cell we quantified 60 features (Bakal et al., 2007). We used the ‘cell area’ feature as a proxy of cell size (Facchetti, Knapp, Chang, et al., 2019). Statistical analysis confirmed the ‘cell area’ distribution means were distinct (N-Way ANOVA, P < 0.05)(**F1A**), demonstrating extensive inter cell line size heterogeneity. We performed an exhaustive series of Wilcoxon rank-sum tests between area distributions (all distributions found to have unique medians, P < 0.001). By retrieving the W-statistic, we calculated the ‘common language effect size’ for each comparison. This produced a matrix of pairwise comparisons between all cell lines that measured the degree of difference in median area between them. Clustering the lines according to this difference let us define three area classes (**F1B**): Class 1; low mean (small cells), low variance high positive skew, class 2; moderate mean, (larger cells) moderate variance, moderate positive skew, and class 3; high mean (largest cells), high variance low skew (**F1C**). BRAFKD/NRAS cells tended to be larger. NRAS and BRAF active cells spanned the range of sizes. Though populations exhibited different extents of variance, we found that across all distributions, the mean cell area linearly scaled with the variance of cell area (R = 0.93), and the coefficient of variation differed only modestly between cell lines (**F1D)**. This suggests that the lines have different size ‘set points’ at which proliferation occurs rather than altered control.

### Cell size relates to DNA content and DNA cytoplasm ratio

Previous studies have indicated that DNA content (Amodeo & Skotheim, 2016), and concentration (Neurohr et al., 2019), are major determinants of cell size. However, Flow-Assisted Cell Sorting (FACS) analysis revealed that ploidy was weakly associated with cell size across lines (**SF2**) For example, both small (4434 (460um^2), 5555 (490um^2)) and large (B14341 (1500um^2), 17864A (900um^2)) cell lines are largely 2N; all exhibited partial 4N populations. We note that 21917 (800um^2) and 24038 (2400um^2) were almost entirely tetraploid.

To further examine the relationship between DNA content and size in single cells, we then quantified the nuclear content (as judged by integrated Hoechst intensity) across lines. This metric differs from ploidy because it considers the amount of Hoechst staining within the nucleus, which can be affected by factors such a packing and aneuploidy in addition to polyploidy. We observed a linear relationship between nuclear content and cell area across lines both between (F1F) and within lines (F1G). We then investigated how the DNA/cytoplasm ratio (D/C) scaled with cell size and identified two distinct clusters of cell lines; A set of smaller cell lines with a relatively high D/C, and large set with a relatively low D/C ratio (F1H). Thus, cell size in these cell lines is related to DNA content and concentration, but ploidy plays a comparatively minor role.

### Translation throttles CCND1 accumulation in response to upstream signalling

To understand the drivers of size in our cell lines, we constructed a proteomic dataset capturing 9,215 total peptides, identifying phosphorylation events on 4,312 peptides, with a total of 2,1355 unique phosphorylation events detected (**SD2**). Peptide expressions, normalised to reflect the relative difference in mass fractions across cells lines (Methods), were correlated to cell areas revealing proteins whose concentrations continuously scale with size.

Previous studies have demonstrated that in normal cells there is a critical RB1 concentration at which cells commit to division (Zatulovskiy et al., 2020). In normal cells, size dilutes RB1 with respect to a constant level of CCDN1 to drive proliferation. However, whether the concentration of RB1 and CCND1 determines a similar set point in cancer cells, especially in melanoma where mitogenic signalling can drive CCND1 translation is unclear (Gennaro et al., 2018; Vízkeleti et al., 2012). As a starting point, we thought to investigate expression of RB1 and CCND1 in our lines. Notably 10/11 of the studied cell lines express detectable RB1, and mean RB1 levels strongly sub-scale with size (**F2A**). Specifically, larger lines exhibited lower mean concentration of RB1. Thus, as within lines (Zatulovskiy et al., 2020), RB1 sub-scales with size between lines.

Notably, CCND1 abundance was also found to broadly negatively correlate/sub-scale with size (F2B). This suggests differential regulation of CCND1 levels between lines. We do note that two lines, 21015 BRAF and C876 NRAS, two of the largest cell lines investigated, exhibited a stark upregulation of CCND1 beyond that would be predicted based on size and RB1 levels. (**F2B**). However, the ratio of RB1 to CCND1 is largely invariant with size (**SD2**). We conclude that the set point, where CDK4/6:CCND1 activation exceeds RB1 concentration to drive proliferation is thus similar across lines.

To understand the molecular basis for the inter-line scaling of CCDN1 we established a method to quantify the signalling activity upstream of CCND1; utilising the phosphorylation state of transcriptional regulators of CCND1, as defined by the ENCODE database ((Dunham et al., 2012)), (henceforth labelled CCND1regs) across different melanoma lines. All phosphorylations used in the analysis with known causative kinases or documented cellular effects (as determined via the Phosphositeplus ((Hornbeck et al., 2015)) database) are detailed in **SD4**. These include several canonical upstream regulators of CCND1 transcription such as BRAF-MEK-ERK as well as JUN and MYC amongst others. Across lines we found that the phosphorylation of the majority of CCND1regs follow a similar trend to RB1 (or ∼CCND1) expression, negatively correlating with size (**F2D**). These pathways were largely upregulated in small cells (Class I), consistent with the presence of activating mutations in BRAF and NRAS (**table 1**). These pathways were also downregulated in large cells (Class 2/3) consistent with the fact that some of these lines have inactivating mutations in BRAF which inactive kinase activity (Bhargava et al., 2016; Malin Pedersen et al., 2014).

Interestingly, despite both negatively correlating with cell size, we observed a negative relationship between cell size and the ratio between RB1 and CCND1reg expressions, showing that larger cell lines in fact exhibit more pro-CCND1 signalling per molecule of RB1 than smaller cell lines. Moreover, this shows that that the low levels of CCND1 in larger lines are not due to reduced mitogen signalling alone.

Investigating this phenomenon, we identified proteins whose expression correlated with the RB1/ pCCND1reg ratio (**F2E**) and conducted SAFE analysis (**F2F**). We observed that low RB1/ pCCND1reg ratio (i.e. in large cells) is associated with lower expression of ribosomal and spliceosome proteins (e.g. RPL26, SNRPE) (**F2G, SD7)**. Thus, as these cells have less CCND1 than would be expected, this suggests that reduced biosynthesis inhibits the conversion of CCND1 signalling to functional CCND1.

Taken together, these data suggest that in large cells, while upstream activity of CCND1 regulators is high relative to RB1, decreased translational efficiency “throttles” CCND1 protein accumulation.

### Proteome wide identification of sub- and super-scaling factors

We next sought to describe more broad differences in protein expression between the lines. By plotting correlations of protein mass fractions with size versus the fold change across big and small cell lines, we conducted a volcano analysis (**F3A)**. We classified size-correlated peptides as ‘Hits’ (Fc > 1.33 or < 0.66, P < 0.05), which were then sorted into two groups. One group of peptides are those expressed in class I (small) cells (thus, sub-scaling) and those expressed in class2/3 (large) cells (super-scaling). We conducted an ontological analysis on all the hit peptides (**F3B**). Small cells were enriched for sub-scaling proteins encoding regulators of cell cycle and mitotic processes (labels including: ‘Cell cycle process’, ‘mitotic cell cycle process’, ‘cell division’, P < 10^-9). These proteins included checkpoint kinases ATM, BRCA1, and WEE1; and the mitotic cyclin CCNB2. In contrast, class 2/3 large lines were statistically enriched for super-scaling peptides from lipid/glycolipid metabolic processes, and components of the extra-cellular matrix (ECM). These included: MVK, MVD, ACAT2 and COL2A1 (all ontology enrichments significant to at least P < 10 ^-3). (**SD3/4**) (**F3E for examples**).

To capture how protein kinase activity may also sub- or super-scale, we analysed the set of proteins for which at least one phosphorylation was detected; using the same system as that above (**F3C/D**). In Class I small cells we observed a clear enrichment of sub-scaling phosphopeptides from cell cycle regulators (‘mitotic cell cycle’, ‘cell cycle process’, ‘cell cycle’, P < 10^-9, eg: BRCA1, SPDL1, LIG1) and biosynthetic processes (‘regulation of cellular biosynthetic process’, ‘regulation of macromolecule biosynthetic process’, ‘positive regulation of RNA metabolic process’, P < 10^-9, eg: mTOR, TSC1, EIF4B, EIF4G1, MED26) including the canonical mTORC1 activating phosphorylation, S2448 (Chiang & Abraham, 2005) implying upregulation in small cell lines. Class 2/3 larger cell lines were enriched for super-scaling phosphorylations on GTPase and cytoskeletal regulators (‘positive regulation of GTPase activity’, ‘cell junction assembly’, ‘regulation of cytoskeleton organization’, P < 10^-4, e.g: CTNNB1, LATS1, ROCK1, ARHGEF5, ARFGEF1, ARHGAP12) and also set of growth regulators (‘regulation of macromolecule biosynthetic process’, P < 10^-6, PDGRFA, PDGFRB, IRS1, PRKCD, DEPTOR) implying differential regulation of biosynthesis across the size range (**SD3, F3E/F/G for examples**). These included activating phosphorylations on ROCK1 (S1341) and a CTNNB1 degradation signal (S29) (Hornbeck et al., 2015). Notably, DEPTOR phosphorylation and expression super-scaled with size implying downregulation of insulin-TOR signalling; this is consistent with the observed reduction in mTOR S2448 expression. See **SF4/5** for a full set of mTOR and cytoskeletal phosphorylations associated with cell size.

We then sought to define a regulatory network of proteins that scale with size. By integrating protein-protein interaction data (Franceschini et al., 2013) (see **SD2** for these unfiltered hits) with our list of phospho-peptide and peptide whose mass fractions correlated with size, we derived a protein-protein network (Shannon et al., 2003) (**F3H**). At this stage, we replaced scaled abundance of the phosphopeptide with an ‘adjusted abundance’, measuring the phosphopeptide abundance relative to the amount of peptide detected (Methods). This way we could reveal peptides that were more/less phosphorylated than expected, given their mass fraction.

Application of the SAFE algorithm (Baryshnikova, 2016) revealed networks of super- and sub-sclaing proteins. These networks were enriched for ontological themes of proteins/phospho-proteins with high expression in each class (**F3H**) ((Ashburner et al., 2000; Carbon et al., 2021; Eden et al., 2009; Mi et al., 2019)). Analysis of these networks echoed the prior results, that it is the disproportionate expression and phosphorylation of ‘G2/M control’ phosphopeptides that defines our smaller cell lines (e.g. BRCA1, WEE1, CCNB2, ATM), including BRCA1 S686 an AKT1 target and stabilising phosphorylation (Hornbeck et al., 2015). In larger lines we observed altered phosphorylation of ‘translation control’ (e.g. eEIF4B, EIF5B), ‘spliceosome machinery’ (e.g. SNRPE, SF3B4), ‘cell adhesion’ peptides (e.g. YAP1, YES1, CTNND1) and increased expression of ‘growth signalling’ (e.g. EGFR, PDGFR) peptides (**F3H**). Interestingly, despite mTOR S2448 sub-scaling, many phosphorylations enriched in larger cell lines correspond to insulin signalling events such as EIF4B S497 (Hornbeck et al., 2015) implying differential interpretation of upstream signals in big/small lines.

### Inflammatory transcripts enrich in larger cell lines

We next performed transcriptomic experiments to gain further insight into the relationship between size and signalling network organization and activity. We measured the abundance of 24988 RNA molecules overlapping with 9290 measured peptides (**SD5**).

We conducted a volcano analysis, as described prior (Methods), to identify a list of transcripts that super- and sub-scale with size across lines (**F4A**). mRNA’s relating to cell cycle regulation (‘regulation of cell cycle’; BARD1, WEE1, E2F8, RB1) and control of gene expression (‘negative regulation of gene expression’, ‘chromatin organisation’, ‘cell differentiation’; e.g: SIN3A, SOX2, HMGA1) sub-scaled with cell size **(F4B/C) (SD3)**. These observations are in line with observations that small cells express relatively higher levels of cell cycle regulatory proteins such as WEE1. Transcripts pertaining to inflammation processes (‘inflammatory signalling’, ‘Interferon signalling’ including: STAT1, IFIT1, IRF5 (and 7) and ADAR) were upregulated in larger cell lines.

In conjunction with that observed in the proteomic data, these data show that compared to Class I smaller cells, Class 2/3 larger cells have decreased levels G2/M regulators, altered metabolism and biosynthesis, and increased expression of inflammatory effectors.

### Transcription regulates ribosomal scaling

To examine the role of translation and transcription in size and proliferation control, we first related mRNA and peptide abundances in each line. Correlation coefficients between mRNA and expression ranged between 0.56-0.38, in agreement with previous studies (Gry et al., 2009) (**F5A/B**). We then calculated correlations at the gene level, across cell lines. Strikingly, this revealed that for most genes there is poor correlation between mRNA fraction (reads of gene/total reads in the cell line) and peptide mass fraction. Indeed, of those that exhibited significant correlations (1116/9290), many showed negative relationships (277/1116) (**F5C**).

By conducting ontological analysis on the genes with significant, (P < 0.05, R > 0.55, n = 11) positive correlation with peptide abundances, we observed an obvious enrichment of cell cycle and DNA repair/replication genes (‘cell cycle process’, ‘DNA Replication’, ‘mitotic cell cycle phase transition’, P < 10 ^-6, eg: ‘BRCA1’, ‘CCNB2’, ‘CCND2’, ‘CCNA2’, ‘BRIP1’, ‘CDK4’, ‘ECT2’) (**F5D/E**). Interestingly, this suggests that the sub-scaling of G2/M regulators with size is occurring at the transcriptional level; as cells get larger fewer G2/M transcripts are being produced reducing peptide expression.

To identify genes that were strongly correlated in specific size classes we split the transcript/proteomic datasets into ‘large’ and ‘small’ subsets, comprised of cell lines with sizes above/below the mean (900um^2), and recalculated the correlation coefficients between mRNA and peptide fractions. At the gene level, In the smaller subset, we observe enrichment of inflammatory, adhesion and cell cycle regulators (‘type 1 interferon signalling pathway’, ‘regulation of cell adhesion’, ‘mitotic cell cycle checkpoint’ P < 10^-5, eg: ‘IRF9’, ‘IFIT3’, ‘STAT1’, RHOD’, ‘CTHRC1’, ‘CCNB2’, ‘BRCA1’) (**F5F-H**). In large cells, there was a strong correlation between mRNA and protein mass fraction of ribosomal and translational genes (n = 5, P<0.05, |R| > 0.75) (‘translation’, ‘cytoplasmic translation’, ‘cytoplasmic large ribosomal subunit’, P<10^-9, eg; RPL26, RPL8, RPL23, RPL5, ETF1) (**F5I-K**). This suggests, like that observed for G2/M regulators, that control of the expression of translational components occurs through transcription, rather than translation, at larger cell sizes. Indeed, correlating RNA pol1-3 component expressions (**F5L**) to the mRNA abundance of ribosomal components, we note a clear positive relationship (for POL1/2). This extended to the peptide abundance suggesting that the production of ribosomal peptides is transcriptionally limited in these cell lines. This was not observed in smaller cell lines; whilst the correlation between pol1/2 expression and peptide expression was maintained, the relationship with mRNA was disrupted indicting a translational dependency in smaller lines (**SD5/3**).

### Validation of size scaling relationships in an independent panel of melanoma cell lines

To assess the universality of our size-scaling relationships, we extended our panel of 11 lines to include a further 12 comprised of the same genotypes as before and conducted further quantitative morphological analyses and phospho-proteomic experiments and repeated the volcano analysis. In contrast to the previous analysis, we note an additional ‘arm’ of the volcano plot indicating a subset of peptides extremely enriched in larger cell lines (**SF3**). We hypothesise that tis result represents gene overexpression rather than super-scaling relationships. Initially including these genes in the analysis, we found that upregulation of apoptotic effectors (‘apoptotic signalling pathway’, ‘positive regulation of mitochondrial membrane permeability involved in apoptotic process’, ‘necrotic cell death’, P < 10^-3, eg. ‘BOK’, ‘TLR3’, ‘TLR4’, ‘BCL2’, ‘TICAM1’), and lipid/carbohydrate metabolism (‘lipid metabolic process’, ‘small molecule metabolic process’, ‘oligosaccharide metabolic process’, P < 10-8, eg. GAA, NEU1, ALG11, ACOX3) is most associated with large cell lines. Excluding the ‘overexpressed’ genes, we observe enrichment of lipid metabolic proteins alone (‘lipid biosynthetic process’,’ lipid metabolic process’, ‘sterol metabolic process’, P < 10^-5). In contrast, we observe a clear enrichment of cell cycle (‘cell cycle process’, ‘cell cycle checkpoint’, P < 10^ -15, eg. CDK2, CCNB2, CCNA2, CDC45), mitotic (‘mitotic cell cycle checkpoint’, ‘chromosome segregation’, P < 10^-10, eg. SPDL1, ECT2, PLK1) and DNA repair (‘DNA repair’, ‘cellular response to DNA damage’ P < 10^-10, eg. BRCA1, LIG1) peptides in smaller lines indicating sub-scaling (**SF4**) (**SD3**).

Enacting the same analysis for the phospho-peptides (**SF4**), we again observe additional ‘arms’ indicating gene overexpression in big/small cell lines. We first calculated enrichments for the two ‘central arms’ finding phosphorylations on cell cycle, DNA repair, and biosynthetic regulatory peptides (‘cell cycle process’, ‘DNA repair’, ‘regulation of macromolecule biosynthetic process’, P < 10^-7, eg: ‘BRCA1’, ‘CHEK1’, ‘CDK4’, ‘EIF4B’, ‘RPS5, ‘EIF3G’) enrich in smaller cell lines. In larger cell lines, phosphorylations pertaining to cytoskeletal and growth factor signalling (‘Regulation of GTPase activity’, ‘cytoskeletal organisation’, ‘cell adhesion’, ‘regulation of epidermal growth factor receptor signalling pathway’, P < 10 ^-3, eg; ARHGEF6, GIT1, TSC2, ROCK1/2, CDC42, TLN1, AKT1/3) are enriched. Within each overexpressed group, we found that larger cell lines were upregulating GTPase signalling elements (‘positive regulation of GTPase activity’, ‘regulation of small GTPase mediated signalling’, ‘Rho protein signal transduction’, (P < 10^-10, 10^-10, 10^-5 respectively), eg. ARFGAP1, TIAM2, ARHGAP1) whilst upregulations in smaller cell lines followed no theme (**SF3, see SF3 for examples**) (**SD3**). We then investigated which ontological themes were enriched in both analyses finding good agreement, a full discussion of this analysis can be found in the supplemental information. Interestingly we recover a large, BRCA1 centric set of interacting genes in both analyses, implicating BRCA1 in size-dependent phenomena (**SF3**).

These data corroborate our previous analysis, strengthening the claim that G2/M and DNA repair processes define smaller melanoma cell lines, (with associated peptides sub-scaling with cell size), whilst cytoskeletal organisation and the rewiring of lipid metabolism define larger cell lines (peptides super-scaling with size).

### Cell growth rate scales with cell size despite downregulation of biosynthetic effectors

Having observed sub-scaling of ribosomal and spliceosome peptides expression, differential phosphorylation of biosynthetic regulators, depressed proliferative signalling and a clear inflammatory response in larger cell lines, we expected them to exhibit a notably decreased growth rate. To investigate this, we live imaged two cell lines from each genotype spread across the observed range of cell sizes and quantified the average rate of growth as the area gain per time, (um^2/hour). To our surprise, growth rate was found to increase with cell size despite the observed downregulation of biosynthetic effectors (**F6A**) and proliferation rate was only modestly affected (**SF3**). We note, however, that this relationship does not appear linear suggesting that the system behind this phenomenon begins to fail at large cell sizes. These data show that larger melanoma cell lines can maintain cell growth without the scaling of classical growth regulators.

Investigating mTOR signalling specifically, we note that while the primary activating mTOR phosphorylation sites (S2448, S2481; phosphorylations responsible for signalling through mTORC1/ 2 respectively (Chiang & Abraham, 2005)) are under-phosphorylated in larger cells (phosphopeptide abundance is lower than expected given peptide abundance), many upstream regulators exhibit phosphorylations typical of insulin driven RTK signalling. However, these genes were differentially phosphorylated across the cell sizes; for example, IRS1 S414 is enriched in smaller cell lines whilst IRS1 T448 is enriched in larger lines, despite both being related to insulin signalling (Hornbeck et al., 2015). These data further indicate that differential, rather than reduced, RTK signalling across sizes leads to the observed downregulation of biosynthetic effectors in larger cell lines (**SF5**).

Given the altered signalling state and having noted an upregulation of cytoskeletal peptide expression and phosphorylation in larger cell lines, we were interested in the state of canonical RTK driven pathways of cytoskeletal activation. Interestingly, we noted that HER2, (1108), SRC (S17), PAK4 (S476), ROCK1 (S1341), VASP (S317) and LIMK1 (S298), (all of which activating phosphorylations (Hornbeck et al., 2015)) amongst others, where disproportionately abundant in larger cell lines indicating that RTK-driven cytoskeletal activity is upregulated (**SF6**). We hypothesise that larger cell lines have skewed their RTK-signalling machinery toward the activation of cytoskeletal rather than anabolic processes. This may lead to increased spreading of the cell.

### Theoretical modelling suggests decoupling of low mitogen and growth signalling drive proliferation of larger cells

Noticing that growth rate is maintained in larger cell lines in the background of downregulated proliferative and biosynthetic signalling, we sought to understand the significance of this effect at a more systems level. We utilised a simplification of recent models, where the transition rate between cell cycle stages is governed by a power-law relationship with cell size:

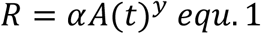

Following previous studies (Nieto et al., 2020), the transition time probability distribution under an exponential growth condition is given as:

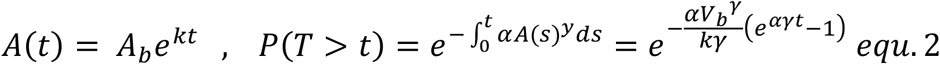

Indicating that the *γ*^th^ power of the added area follows an exponential distribution centred on *γk/α*;

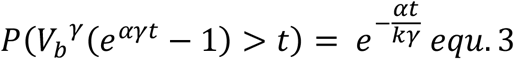

Taking y =1, the mean added mass equals *k/α*. We could capture similar behaviour when considering the simpler case where the probability of transitioning between cycle stages and growth rate are taken to be a constant within a cycle but are adjusted according to cell division size (Adiv), (P = αA_div_, β = kA_div_, respectively) defining a Poissionian system. We believe that this simplification provides a useful tool for the understanding of cell size determination in the adder case. Here we have assumed adder-like behaviour, as small errors in sizer mechanisms can lead to phenomenological adder systems (Facchetti, Knapp, Chang, et al., 2019). Using these, we could derive expressions for the expected proliferation rate and added size, given as exponential distributions;

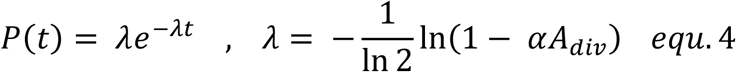

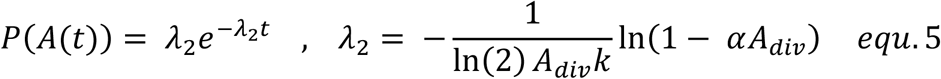

We include the derivations in the supplemental information. This facilitated the construction of a simple system of equations dictating cell size:

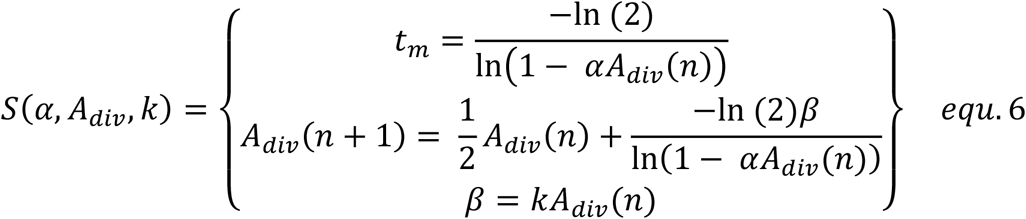

Where ‘tm’ is the mean proliferation time, and ‘n’ is the number of proliferative cycles that have passed. Perturbing the parameters of this system, we find it is stable to perturbations in A_div_ (**F6B**) but unstable to changes in α or k. That is, a constant mean size is maintained under this system that may be adaptively regulated by modification of α, related to mitogenic signalling and k, controlling growth rate.

If alpha is perturbed, the proliferation rate initially decreases, but exponentially decays back to the initial value across successive division cycles. Thus, if the alpha and k parameters are independent, cell growth provides a means to ‘correct’ proliferation rate under perturbation to mitogenic signalling. (**F6B**)

Using equations 4 and 5, we could derive the moments of the expected size distribution (**SI**). This is a hypo-exponential function, with a mean and variance given as:

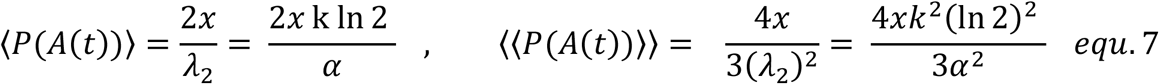

Where ‘x’ is the number of ‘stages’ in the model of cycle. This yields a coefficient of variation;

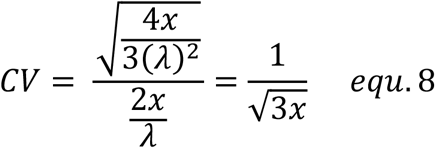

The cell lines have CV’s of ∼ 0.7 – 0.6, and x was calculated to range between 0.7 and 0.9. For simplicity, we took x = 1 from this to avoid complications stemming from a decimal number of cell cycle stages (although this may be rationalised as the cell ‘skipping’ cell cycle regulation every 1/x divisions). We note this approach is only feasible when the CV > ∼0.25 as the differences in CV values for neighbouring ‘x’ tend to 0 as x increases; this is equivalent to an ∼ 5 stage system. (**F6C**)

These results allow us to define a simple and efficient algorithm to calculate predicted cell size distributions (methods) (**F6E**). ‘α’ values were fit to experimentally determined area distributions by minimising the Kullbeck-Leibler divergence (Andrew, 2004) between measured and calculated distributions (beta is a measured parameter); ‘*α*’ values decrease with increasing cell size (**F6F**) (**Table 2**). The simulation accurately recapitulated much of the measured data, however, In the case of larger cell lines, the model partially under-predicted the abundance of ‘small’ (A < 500um^2) cells (see 24038 (2500um^2), C876 (1600um^2), 17864A (1000um^2) and B14341 (1500um^2) (**F6G**). A summary of the model can be found in **F6H**.

**Table 2:**
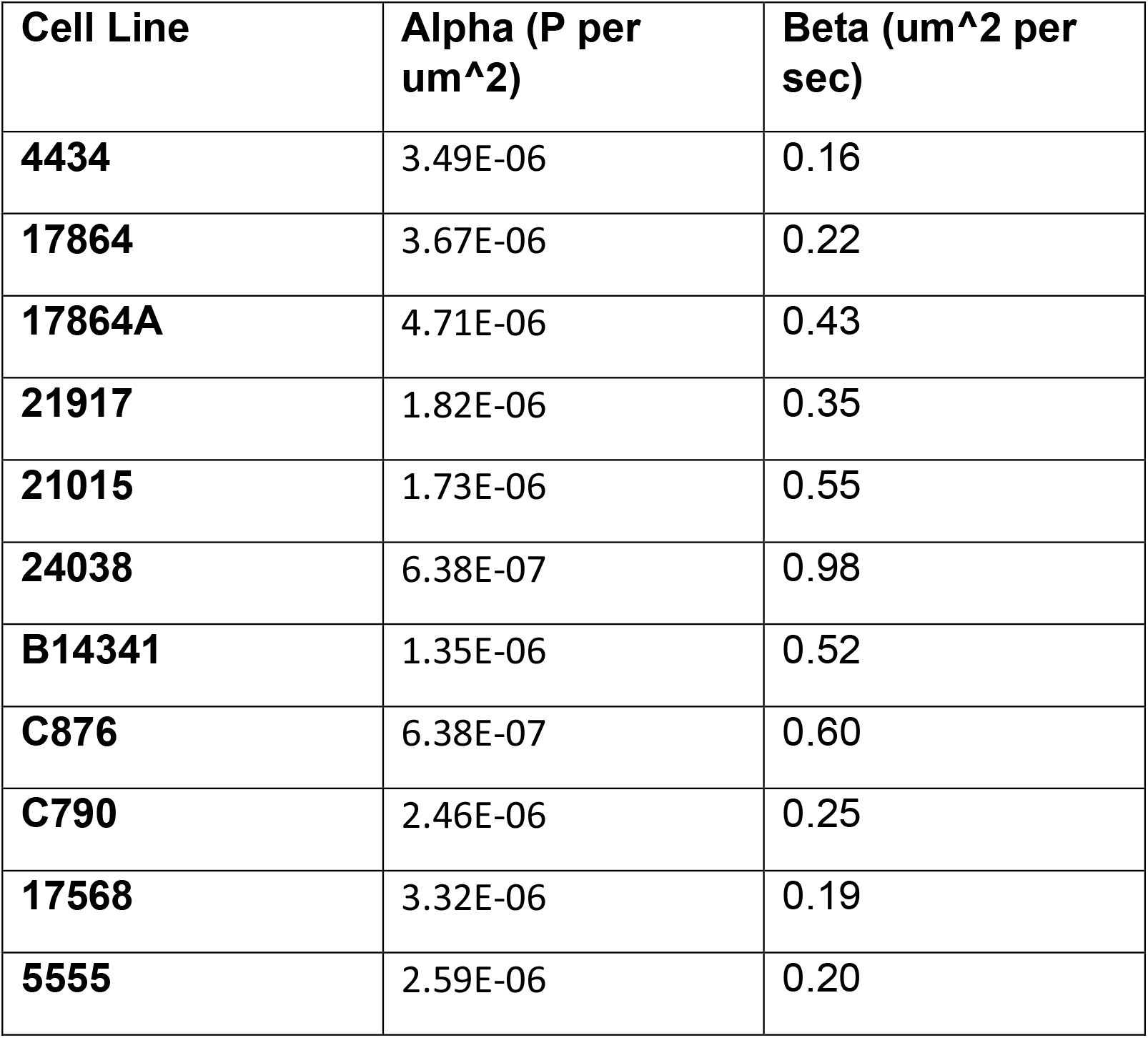
Model Parameter Values.

| **Cell Line** | **Alpha (P per um<sup>2</sup>)** | **Beta (um<sup>2</sup> per sec)** |
| --- | --- | --- |
| **4434** | 3.49E-06 | 0.16 |
| **17864** | 3.67E-06 | 0.22 |
| **17864A** | 4.71E-06 | 0.43 |
| **21917** | 1.82E-06 | 0.35 |
| **21015** | 1.73E-06 | 0.55 |
| **24038** | 6.38E-07 | 0.98 |
| **B14341** | 1.35E-06 | 0.52 |
| **C876** | 6.38E-07 | 0.60 |
| **C790** | 2.46E-06 | 0.25 |
| **17568** | 3.32E-06 | 0.19 |
| **5555** | 2.59E-06 | 0.20 |

Taken together our modelling has shown that given a proportionality between cell size and division probability, increased cell growth is an effective means of triggering proliferation when scaling of proliferative factors is perturbed, for example, by a reduction in mitogenic signalling.

## Discussion

We have identified scaling relationships between cell size and peptide/gene expression in melanoma. Expression and phosphorylation of G2/M, DNA-associated and biosynthetic peptides exhibited a clear sub-scaling relationship with cell size across two independent panels of melanoma cell lines, whilst expression of lipid metabolic genes and phosphorylation of cytoskeletal regulators showed the reverse. This is in strong agreement with numerous recent studies investigating the relationships between cell size and gene/peptide expression; identifying histones as sub-scaling components (Amodeo et al., 2015), observing an upregulation of lipid metabolism in larger cell lines (Neurohr et al., 2019), noting a decreased abundance of translational components and translation rate in large polyploid cells (Yahya et al., 2021) and a full proteome survey of scaling components in human lung fibroblasts (Lanz et al., 2021).

Interestingly, we observe that the mean RB1 mass fraction decreased with increasing cell size corroborating the findings of recent studies associating RB1 (and Whi5) dilution to size determination and control (Schmoller et al., 2015) (Zatulovskiy et al., 2020). This trend extended to the abundance of phosphopeptides associated with CCND1 transcription and the abundance of core ribosomal and spliceosomal peptides. These data suggest that the state of the RB1-CCND1 axis in melanoma, or indeed an RB1-dilution system (Zatulovskiy et al., 2020), is sensitive to both the strength of proliferative signalling and translational capacity of the cell in melanoma. Reduced signalling and protein production may decrease CCND1 abundance, and therefore, RB1 must dilute further to induce division commitment; thereby delaying proliferation until a greater cell size. Interestingly, recent literature suggests G2-driven synthesis of CCND1 (Min et al., 2020; Stallaert et al., 2021) noticing a tight dependence on cellular translation (Min et al., 2020). Translation and mitogen signalling in the prior G2 may colour events in the subsequent G1. ‘Sub scaling’ of G2/M regulators, such as WEE1 and BRCA1, may relate to smaller cells exhibiting increased expression of CCND1.

We also observed an upregulated inflammatory response and decreased DNA/cytoplasm ratio in larger cell lines; phenomena recently related to the onset of cell senescence (Neurohr et al., 2019). But while larger lines appear morphologically senescent, they are clearly not senescent, as they grow and proliferate at a similar rate to smaller cells. Indeed, this finding is particularly striking given the observed downregulation of canonical pro-biosynthetic phosphorylations (for example in the AKT-mTOR pathway, see **SF5**), given the NRAS activating mutations of larger cell lines (or PTEN null mutation in the case of 21015 BRAF). This shows that larger cell lines maintain high growth rates despite downregulation of anabolic pathways and decreased ribosomal mass fractions, qualities typically associated with decreased biosynthesis (Fingar et al., 2004; Serbanescu et al., 2020).

The mechanism behind this phenomenon is unclear but may relate to mechano-biological processes, given the observed upregulation of ECM components and RAC GTPase signalling in larger cell lines. Indeed, mechanical activation of YAP/TAZ signalling has been observed to facilitate growth/proliferation under MAPK inhibition (Kim et al., 2016; Lin & Bivona, 2016). Furthermore, cell volume has recently been tied to substrate stiffness and adherence, engaging in a feedback system with YAP/TAZ (Gonzalez et al., 2018). Interestingly, several studies suggest that actomyosin contractility during cell spreading can also reduce cell volume through the expulsion of water, concentrating cell constituents (Guo et al., 2017; Venkova et al., 2021; Xie et al., 2018). Large cell lines may activate cytoskeletal signalling to concentrate key biosynthetic regulators and sustain growth.

We constructed a simple theoretical model to demonstrate how continued growth under proliferative stress, could maintain the cells proliferation rate. This relied on the probability of a cell transitioning to the next stage of the cell cycle being proportional to its size (Nieto et al., 2020), for example, via RB1 dilution. This is consistent with the recent observation that cell cycle phase lengths across generations are coupled in cancer cell lines (Chao et al., 2019), here via mother cell size (Min et al., 2020). Interestingly, the same study notes this effect may be unique to cancerous cell lines due to a disproportional abundance of regulators acting at multiple stages of the cell cycle (Chao et al., 2019) in effect ‘simplifying’ regulation. Indeed, through analysis of cell size variation, we found our cell lines were most effectively modelled by a one-(growth) stage cycle, implying the dominance of a small subset of proliferative regulators. This suggests a more central role for the RB1 sub-scaling observed in these cell lines.

Taken together, our data shows that despite sub-scaling relationships between key biosynthetic and proliferative regulators and cell size, and a robust inflammatory response, larger melanoma cell lines exhibit a higher growth rate than smaller lines. Theoretical modelling suggests that proliferation may be maintained under mitogenic inhibition by decoupling growth and proliferative signalling. Oncogenic mutations could facilitate this process and may be associated with cytoskeletal activity.

## Methods

### Cell Culture

Cell lines were maintained in standard culture conditions (DMEM+10% FBS). Passage was carried out using 0.25% trypsin-EDTA (GIBCO) followed by centrifugation (1000 rpm, 4 min) and resuspension in complete medium. Cell counting was performed using Countess automated cell counter with trypan blue exclusion (Thermo).

### Growth Curves

Each cell line was seeded into 3 wells of a 6 well tissue culture plate. Cells were incubated in DMEM media with 10% fetal bovine serum and Primocin antibiotic, at 37 degrees Celsius and 5 % carbon dioxide. Cells were imaged at 4 hour intervals using the Incucyte imaging system. 9 fields of view were imaged from each well. Images were segmented using Ilastik image segmentation software to identify individual cells. The number of cells in each field of view was calculated using CellProfiler. Growth curves were plotted using the ggplot2 library from the R programming language.

### Immunostaining

Samples were fixed in freshly prepared 4% PFA/PBS for 15 minutes. Slides were subsequently permeabilized with 0.25% Triton/PBS for 10 mins and blocked with 0.5% BSA/0.02% glycine/PBS for 30 minutes. Primary antibodies were introduced via the same solution in a 1:1000 dilution and left on for 1 hour. The slides were washed with PBS and the same was carried out for the secondary antibodies (kept in the dark to avoid bleaching). Hoechst stain was added post-secondary (1:500) to stain DNA as was phalloidin to stain actin.

### Image Acquisition and Feature Extraction

Image acquisition was performed using an Opera Cell: Explorer-automated spinning disk confocal microscope. 20 fields of view were imaged in each well. Cell segmentation was performed using Acapella software (PerkinElmer). Nuclei were segmented using the Hoechst channel (405-450) and cell bodies defined by the tubulin signal (568-603). Geometric features measured include: the area of all subcellular regions; the length, width, and elongation (length/ width) of the cell and nucleus, cell and nuclear roundness and nucleus area/cytoplasm area. Texture features were also measured representing the distribution of pixel intensities in a cell or subcellular region. Haralick and Gabor features (Fogel & Sagi, 1989; Haralick et al., 1973) as well as SER (“Saddle/Edge/Ridge”, PerkinElmer) features were measured on the Hoechst and tubulin channels.

### Statistical Analysis of Cell Size

Statistical test were carried out in the MATLAB (math works) environment. Cell area data was ‘acosh’ transformed to induce a normal distribution of areas in each cell line and standardise the variances prior to ANOVA and Mann-Whitney/Wilcoxon tests. Standardisation success was determined using the Shapiro-Wilks normalization test, ensuring the data is normally distributed, and the Bartlett test, to guarantee equal variances across lines.

### FACS Analysis

Cells were trypsinized and harvested into a 15ml falcon tube for cell counting. After centrifuging the falcon at 2400rpm for 5 minutes, the supernatant was discarded and the cells were resuspended in 1mL of 1% FCS in PBS. 3mL ice cold 100% ethanol was added dropwise to the cells while slowly vortexing, and left to fix overnight. The cells were then pelleted by centrifugation for 5 minutes at 2400rpm, and resuspended in 5mL PBS. They were incubated at room temperature for 20 minutes. After centrifuging for 7 minutes at 1200rpm, the pellet was resuspended in 1mL of Propidium Iodide (PI) solution through the cell strainer into a FACS tube. The PI solution was made with 1:100 PI at 5mg/mL and 1:1000 RNAaseA at 10mg/mL in PBS. The cell cycle composition was measured using the BDSAria and the data analysed using FlowJo. For EdU (5-ethynyl-2’-deoxyuridine) incorperation assays, cells were treated with a final concentration of 10uM EdU prior to harvesting. Instead of fixing with ethanol and staining with PI, cells were resuspended in 4% PFA for 15 minutes at room temperature. They were then pelleted by centrifugation and PFA aspirated, followed by a wash. 500uL of the appropriate Thermo Fisher Click-iT™ reaction cocktail was added to each sample and incubated for 30 minutes in the dark, according to the manufacturer’s instructions. Cells were washed once, stained and then transferred via a cell strainer into a FACS tube for analysis as above. Washes used 1% BSA in PBS. Staining used 20ug/mL Hoechst added to 0.1% Triton-X in PBS. If applicable, 10^6^ cells were seeded in Falcon T25 flasks and incubated overnight in media containing the appropriate Aphidicolin concentration (total volume of 4mL) before FACS analysis of DNA content as above.

### Proteomics Sample Preparation

Cell pellets were dissolved in 150 μL lysis buffer containing 1% sodium deoxycholate (SDC), 100mM triethylammonium bicarbonate (TEAB), 10% isopropanol, 50mM NaCl and Halt protease and phosphatase inhibitor cocktail (100X) (Thermo, #78442) on ice with pulsed probe sonication for 15 sec. Samples were boiled at 90 °C for 5 min and sonicated for another 5 sec. Protein concentration was measured with the Quick Start™ Bradford Protein Assay (Bio-Rad) according to manufacturer’s instructions. Aliquots containing 100 μg of protein were reduced with 5 mM tris-2-carboxyethyl phosphine (TCEP) for 1 h at 60 °C and alkylated with 10 mM Iodoacetamide (IAA) for 30 min in dark. Proteins were then digested overnight by adding trypsin at final concentration 75 ng/μL (Pierce). The resultant peptides were labelled with the TMT-11plex reagents (Thermo) according to manufacturer’s instructions and were combined in equal amounts into a single tube. The combined sample was then dried with a centrifugal vacuum concentrator. Two technical replicate TMT batches from the same protein extracts were prepared to assess reproducibility. One TMT batch was fractionated offline with high-pH Reversed-Phase (RP) chromatography using the XBridge C18 column (2.1 × 150 mm, 3.5 μm, Waters) on a Dionex UltiMate 3000 HPLC system. Mobile phase A was 0.1% ammonium hydroxide (v/v) and mobile phase B was acetonitrile, 0.1% ammonium hydroxide (v/v). The TMT labelled peptide mixture was reconstituted in 100 uL mobile phase A and fractionated with a gradient elution method at 0.2 mL/min as follows: for 5 min isocratic at 5% B, for 35 min gradient to 35% B, gradient to 80% B in 5 min, isocratic for 5 min and re-equilibration to 5% B. Fractions were collected every 42 sec and vacuum dried. The second TMT replicate batch was fractionated with the Pierce High pH Reversed-Phase Peptide Fractionation Kit according to manufacturer’s instructions.

### Phosphopeptide enrichment

Peptide fractions from the first TMT batch were reconstituted in 10 μL of 20% isopropanol, 0.5% formic acid binding solution and were loaded on 10 μL of phosphopeptide enrichment IMAC resin (PHOS-Select™ Iron Affinity Gel, Sigma) already washed and conditioned with binding solution in custom made filter tips fitted on Eppendorf tubes caps. After 2 h of binding at room temperature, the resin was washed three times with 40 μL of binding solution at 300 g and the flow-through solutions were collected for total proteome analysis. Phosphopeptides were eluted three times with 70 μL of 40% acetonitrile, 400 mM ammonium hydroxide solution. Eluents and flow-through samples were vacuum dried and stored at -20 °C until the LC-MS analysis.

### LC-MS Analysis

LC-MS analysis was performed on the Dionex UltiMate UHPLC 3000 system coupled with the Orbitrap Lumos Mass Spectrometer (Thermo Scientific). Peptides were loaded to the Acclaim PepMap 100, 100 μm × 2 cm C18, 5 μm, 100 Ȧ trapping column at 10 μL/min flow rate. The sample was then subjected to a gradient elution on the Acclaim PepMap RSLC (75 μm × 50 cm, 2 μm, 100 Å) C18 capillary column at 45 °C. Mobile phase A was 0.1% formic acid and mobile phase B was 80% acetonitrile, 0.1% formic acid. The separation method at flow rate 300 nL/min was as follows: for 90 min (or 150 min for the replicate batch) gradient from 10% to 38% B, for 10 min up to 95% B, for 5 min isocratic at 95% B, re-equilibration to 10% B in 5 min, for 10 min isocratic at 10% B. Precursors between 375-1,500 m/z were selected with mass resolution of 120 K, AGC 4×105 and IT 50 ms with the top speed mode in 3 sec and were isolated for CID fragmentation with quadrupole isolation width 0.7 Th. Collision energy (CE) was 35% with AGC 1×104 and IT 50 ms. MS3 quantification was obtained with HCD fragmentation of the top 5 most abundant CID fragments isolated with Synchronous Precursor Selection (SPS). Quadrupole isolation width was 0.7 Th, CE 65%, AGC 1×105 and 105 ms IT. The HCD MS3 spectra were acquired for the mass range 100-500 with 50K resolution. Targeted precursors were dynamically excluded for further isolation and activation for 45 seconds with 7 ppm mass tolerance. Phosphopeptide samples were analysed with an HCD method at the MS2 level with CE 38%, AGC 1×105 and max IT 105 ms.

### Database search and protein quantification

The SequestHT search engine was used to analyse the acquired mass spectra in Proteome Discoverer 2.2 (Thermo Scientific) for protein identification and quantification. Precursor mass tolerance was 20 ppm and fragment ion mass tolerance was 0.5 Da for the CID and 0.02 Da for the HCD spectra. Spectra were searched for fully tryptic peptides with maximum 2 miss-cleavages. TMT6plex at N-terminus/K and Carbamidomethyl at C were defined as static modifications. Dynamic modifications included oxidation of M and Deamidation of N/Q. Dynamic phosphorylation of S/T/Y was included for the phospho-enriched samples. Peptide confidence was estimated with the Percolator node. Peptide FDR was set at 0.01 and validation was based on q-value and decoy database search. Spectra were searched against reviewed UniProt mouse protein entries. The reporter ion quantifier node included a TMT 11plex quantification method with an integration window tolerance of 15 ppm and integration method based on the most confident centroid peak at the MS3 or MS2 level. Only unique peptides were used for quantification, considering protein groups for peptide uniqueness. Peptides with average reporter signal-to-noise >3 were used for quantification.

### Proteomic Analysis

Peptide abundances were scaled relative to other detected peptides in the sample such that they reflect abundance/total protein mass. The expression of each peptide was correlated to average cell line area to derive a correlation coefficient, R, and an area-weighted fold change was calculated between cells lying above or below the mean according to:

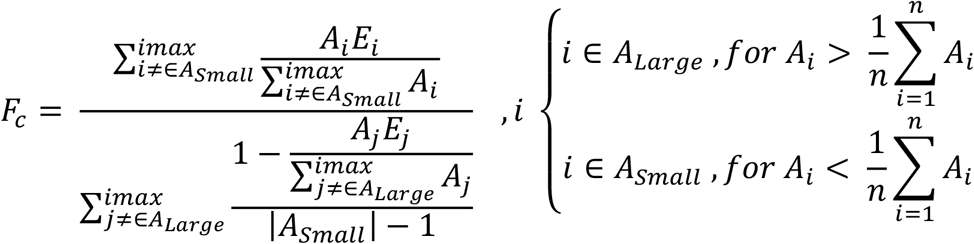

Where Aj is the mean area of the jth cell and Ej is the peptide expression for the jth cell. Lines with areas greater than the mean across lines have their expressions contribute to the large group and vice versa. The weight of this contribution is determined by the lines area contribution to the total of the group. A large contribution results in a higher weighting in large lines and the reverse in smaller lines.

In conjunction with P, the correlation coefficient between peptide and area, the fold change facilitates calculation of a z-score for each peptide, representing its apparent importance to cell area:

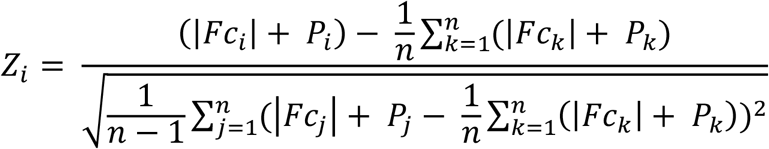

### Network Analysis

High scoring proteins are taken forward and entered into STRING ((Franceschini et al., 2013)) to screen for interactions within the hits. Accepted interactions were those identified experimentally or identified in previous co-expression studies and achieved a confidence value > 0.4. This network was then exported to Cytoscape ((Shannon et al., 2003) for ontological analysis via the SAFE ((Baryshnikova, 2016)) tool. All ‘biological process’ annotations for each node in the network were derived from Geneontology.org’s downloadable database ((Ashburner et al., 2000; Carbon et al., 2021; Eden et al., 2009; Mi et al., 2019)). A binary matrix was constructed; each node (row) would receive either a 1 or 0 in each column (annotation) depending on whether the node was associated with the label. This was then entered into the SAFE Cytoscape plugin, where we used default settings besides a percentile threshold of 10 and minimum neighbourhood size of 5 (**SD5**). A second binary matrix was then constructed now with an annotation set reflecting whether the node was an expression or phosphorylation hit in big or small cells. The same settings were used for SAFE.

### RNA extraction, Quality control and RNA Sequencing

RNA from 11 cell lines was extracted using the RNeasy Mini Kit (Qiagen, #74104) according to the manufacturer’s protocol. The evaluation of the isolated RNA integrity and quantity was carried out by the Agilent TapeStation system using an RNA ScreenTape (Agilent Technologies, #5067-5576).

For the mRNA Library preparation 4000ng of total RNA was treated with TurboDNase to remove genomic DNA contamination, (Invitrogen, #AM1907). PolyA RNA was selected from 1000ng of the purified RNA using NEBNext mRNA magnetic Isolation Module (NEB, #E7490) following manufacturer directions. From the resulting mRNA, Strand-specicific libraries were created using NEBNext Ultra II Directional RNA Library Prep Kit for Illumina (NEB, #E7760). Final libraries were quantified using qPCR and clustered at a Molarity of 300 pM. Sequencing was performed on an Illumina NovaSeq 6000 (Illumina) using PE x100 cycles v1.0 chemistry, to achieve coverage of 25 million reads per sample.

### Transcriptomic Analysis

RNA abundances were normalised across cell lines and filtered through a sigmoidal expression to dampen the effects of extreme over/under expressions warping the analysis. Beyond this, the transcripts abundances were treated identically to the peptide abundances.

### Model Algorithm

The initial cell area distribution is considered a delta function centred on ‘k’/’a’ (the expected mean of the distribution). Every generation, the area distribution is convolved with the mass-gain distribution, computed by performing an inverse Fourier transform on the product of the two distributions respective Fourier transforms. This produces the division area distribution, Ad(A), which must be transformed to Ad(2A) to capture the effects of cell division. We perform this by setting Ab(Ax) = Ad(Ai) + Ad(Ai+1), where ‘I’ = x_n_-x_n-1_ for all x, where Ab denotes the birth size distribution. This is then convolved with the gain distribution as before to generate the next division distribution and so on until a desired number of generations has been reached.

### Numerical Simulation

An initial population of 1000 cells is assigned an ‘alpha’ and ‘beta’ value and a random initial area. At each time step, the division probability for each cell is calculated, according to P = αA_div_, and a random number, ‘r’, is drawn from a flat distribution. Should ‘r’ be less than the division probability of a cell, the cell divides symmetrically in two, adding a new cell to the population with half the size of the mother, and halving the mother size. If ‘r’ is greater than the division probability, the cell size increases according to β = kA_div_. This system continues until a final cell count of 20,000 is achieved.

### Model Fitting Procedure

Initially, alpha values were exhaustively tested (beta is determined from the proliferation and area measurements on a per cell line basis). For each we calculated the Kullbeck-Liebler divergence between the experimental and simulated data **(38)**. For discrete probability distributions defined on the same probability space, X, the Kullback–Leibler divergence from P to Q is ((Andrew, 2004)):

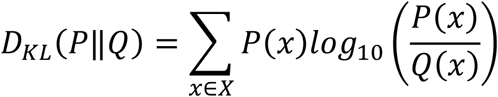

Having identified an approximate-minima from the low resolution parameter screen, we used the values defining this region as an initial state for a stochastic gradient descent minimising along the gradient:

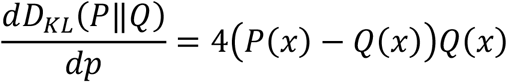

Model fitting was conducted within the commercial MATLAB (Math Works) software’s machine learning toolbox.

## Contributions

I.Jones analysed the data, developed the computational models and wrote the manuscript. T.Higo and L.Dent maintained and imaged the cells and, with H.Shree, conducted the FACs analysis. T. Roumeliotis and J.Choudhary gathered and prepared the proteomic data for analysis and M.A.Garcia and T.Higo the transcriptomic data. M.Pederson developed the cell lines, C.Bakal conceived and designed the research.

## Acknowledgments

We thank Bela Novak and his group for their helpful comments on the early formulations of the theory. This work was supported by a Stand Up to Cancer UK Programme Foundation Award from Cancer Research UK to Chris Bakal (C37275/1A20146).

## Data Availability

Proteomics data is publicly available at the PRIDE database: **Project accession:** PXD028339

A demonstration of the model and corresponding numerical simulation may be found at: <u>IJICR/Model_demo</u> (github.com)

Transcriptomic and imaging data are available on request

## FIGURE LEGENDS

**Figure 1:**
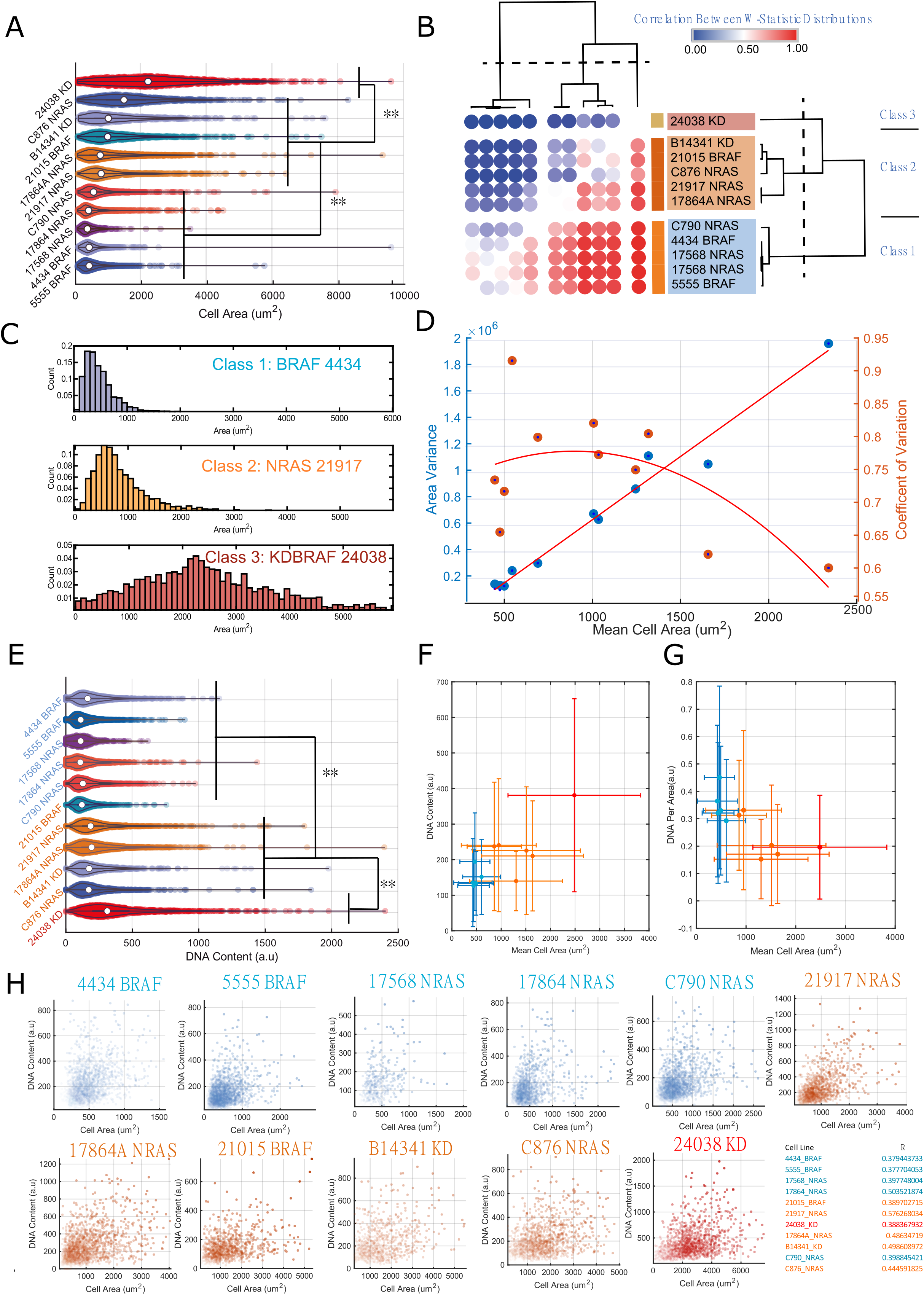
Melanoma cell lines exhibit comparable size control but different cell sizes: A) Violin plot summarizing cell area distributions across lines. Acosh normalised Distribution means were subjected to an 11-way Anova test to confirm the significance of observed differences. ** indicates a P-value < 0.01 B) Heatmap showing a clustering of lines based on effects sizes calculated post Mann-Whitney tests (median uniqueness follows the same pattern as the means P < 0.01). Three distinct area ‘classes’ emerge. C) Sample distributions from each class; from C1 to C3, skew decreases, whilst the means and variances increase. D) Shows the relationship between the means and variances of the area distributions. The mean scales approximately linearly with variance. The coefficient of variation inconsistently varies with cell size. E) As in ‘A’ but for the DNA content distributions. Lines are coloured by size class, and will be throughout the manuscript. F) The relationship between mean cell area and mean DNA content, area positively correlates with cell area. Error bars represent the standard deviation of the single cell data G) Relationship between DNA content and DNA per area. Despite large lines often having more DNA, they exhibit a lower DNA concentration than the smaller cells. H) DNA-content area relationship across all lines at the single cell level. All cell lines exhibit a positive correlation between DNA abundance and size.

**Figure 2:**
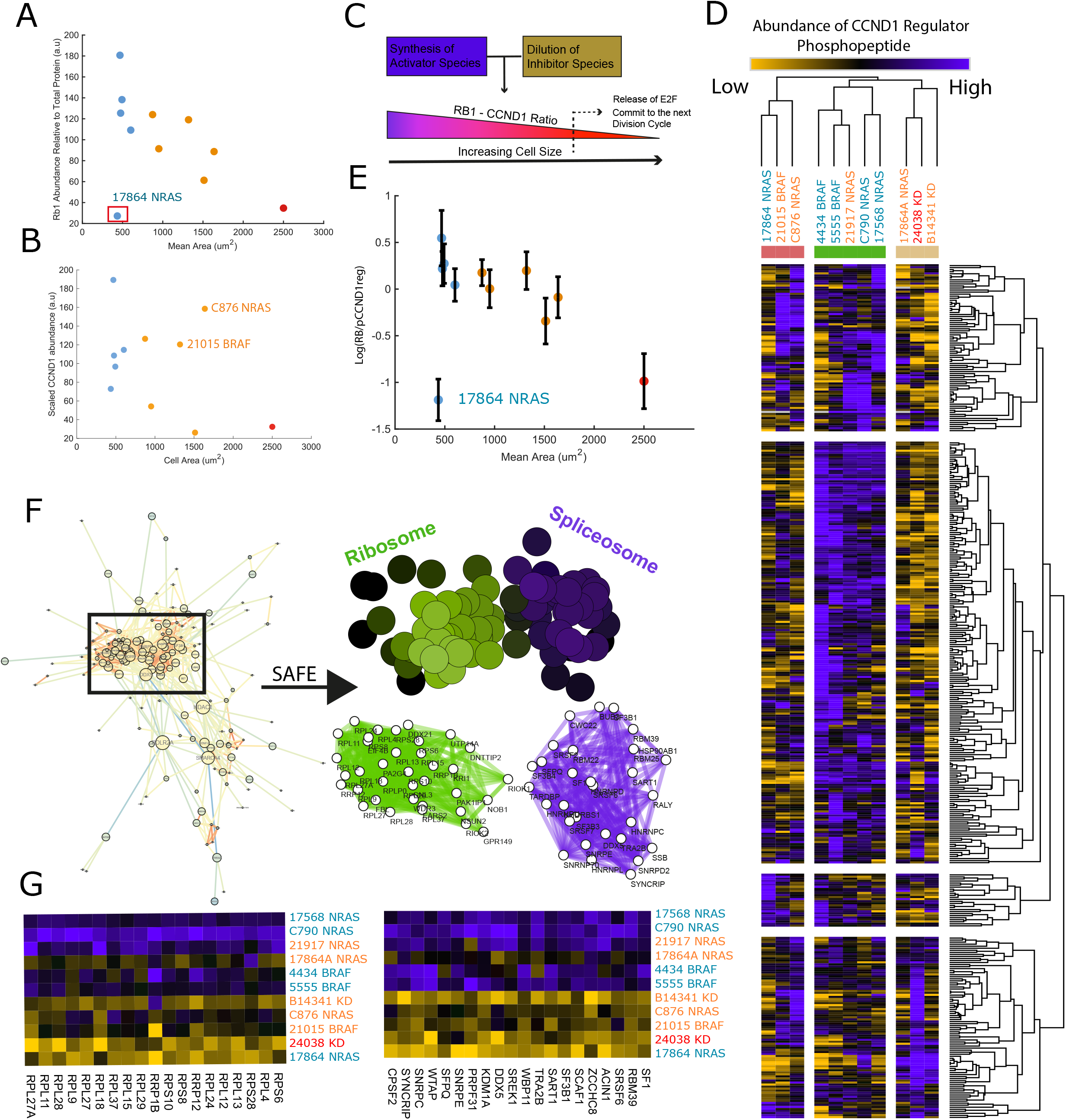
Translation throttles CCND1 accumulation in response to upstream signalling: A) Negative correlation between RB1 mass fraction and cell size. B) Relationship between CCND1 mass fraction and cell size, C876 and 21015 exhibit surprising high levels of CCND1 given their RB1 abundance C) Cartoon schematic depicting the role of RB1 in the dilution model of G1/S transition. D) heatmap depicting the ratio of RB1 against detected CCND1 regulator phosphopeptides, ratios are typically lower in larger lines. E) The average value for all RB1/rCCND1 ratios for each line plotted against cell area. F) Network describing interactions between proteins correlating with RB1/pCCND1reg. SAFE overlay on ‘F’ screening for graph regions enriched for ontological labels. Colour intensity denotes the confidence of the enrichment. G) heatmaps showing the expression of peptides found in enriched regions of the interaction network across lines.

**Figure 3:**
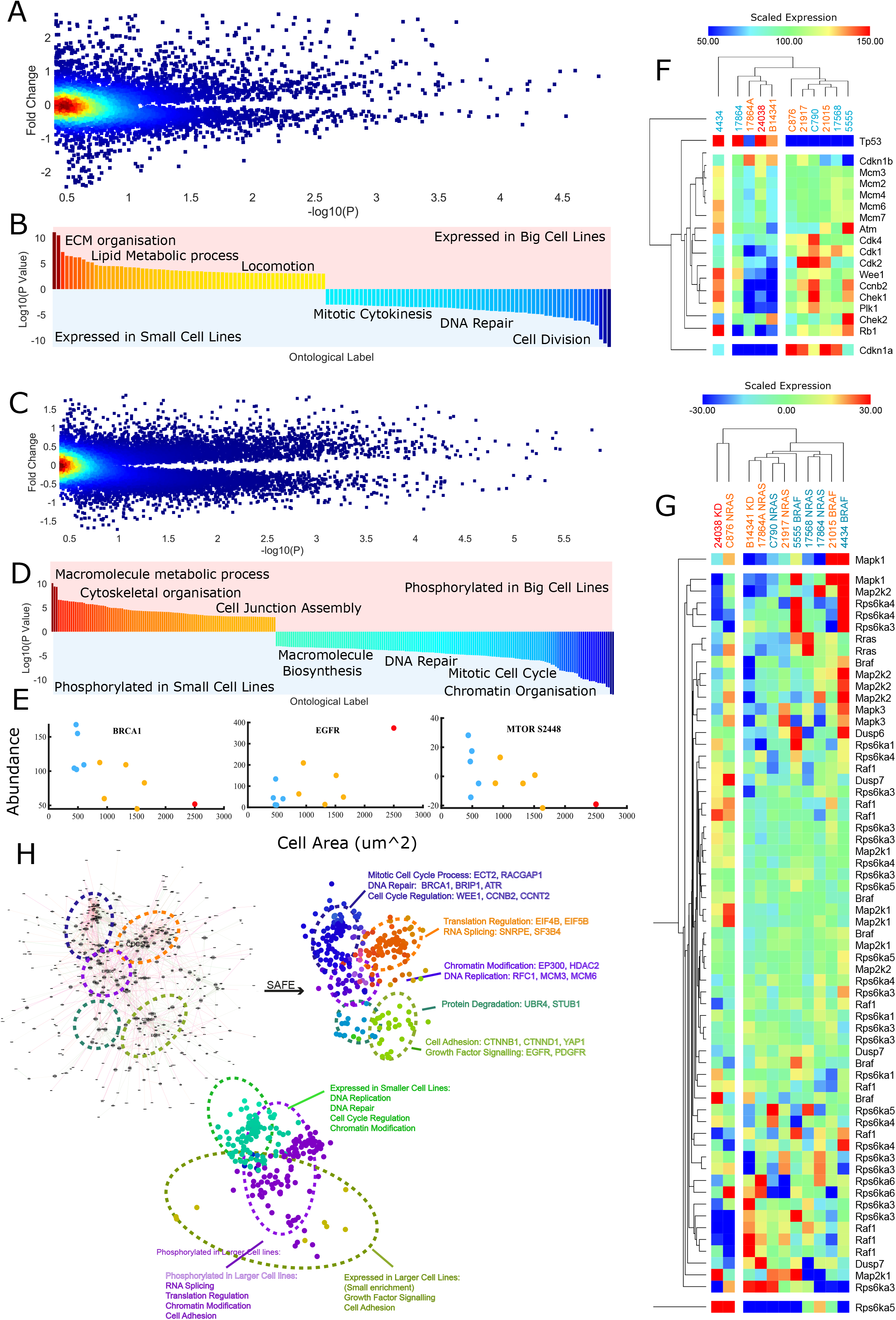
Proteome wide identification of sub- and super-scaling factors. A) Fold change in peptide abundance across large and small cell lines plotted against the significance of the expressions correlation with size (genes achieving abs(log2(fc)) > 0.5, P < 0.05 are taken forward for ontological analysis) Colour represents data point density B) Ontologies enriched in peptides differentially expressed across big/small lines C) Fold change in phosphopeptide abundance across large and small cell lines plotted against the significance of the expressions correlation with size (genes achieving abs(log2(fc)) > 0.5, P < 0.05 are taken forward for ontological analysis). Colour represents datapoint density D) Ontologies enriched in phosphopeptides differentially expressed across big/small lines F) Heatmap of select G2/M controllers revealed to enrich in smaller cell lines. G) Expression of phosphopeptides pertaining to the MAPK pathway H Network derived from screening for interactions within the list of size predicting, kinase regulated, peptides. Interaction data was obtained from the STRING database. Right; SAFE overlay on ‘D’ screening for graph regions enriched for ontological labels. Lower; SAFE overlay on ‘D’ screening for regions with high expression/phosphorylation in large/small cell lines.

**Figure 4:**
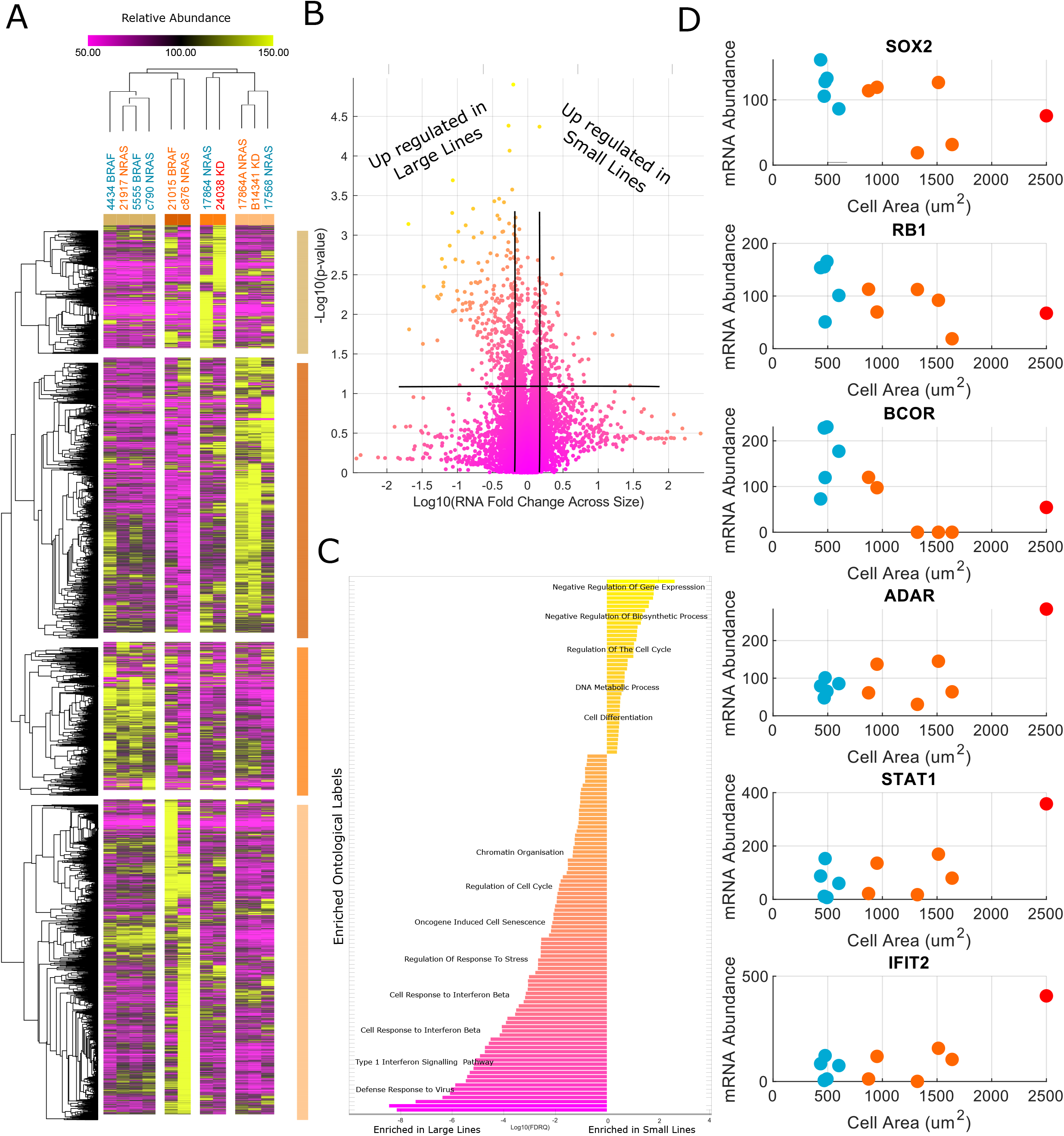
Inflammatory transcripts enrich in larger cell lines: A) Heatmap of transcriptomes across cell lines. Lines/transcripts are grouped through hierarchical clustering conducted using the ‘Morpheus’ software (Broad Institute). B) Volcano plot for the fold change of transcripts across size groups against the RNA-size correlation. C) Ontologies enriched in large/small cell lines, Inflammatory transcripts enrich in larger lines whilst those related to cell cycle and gene regulation enrich in smaller lines. D) Examples of correlating transcripts in either group.

**Figure 5:**
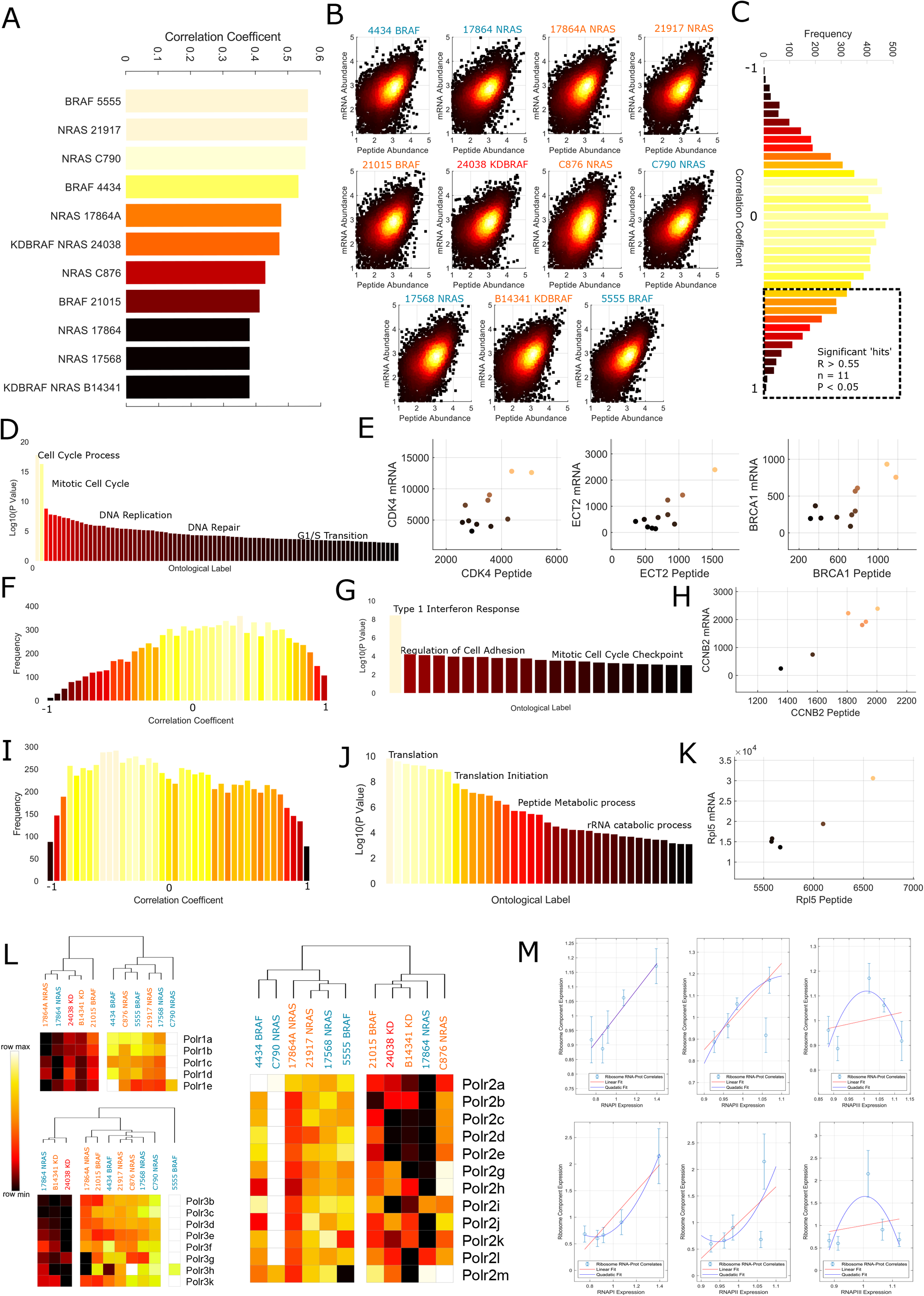
Transcription regulates ribosomal scaling. A) Correlation between gene and peptide expression within each cell line, coefficients range between 0.4 and 0.6 B) Log-log plots of RNA against peptide abundance, colour intensity is proportional to the density of the data points C) Distribution of correlation coefficients between peptide and mRNA abundances across cell lines, dotted box indicates genes with significant correlations. D) Enriched gene ontologies detected in the genes with significant RNA-peptide correlations. E) Example protein-mRNA correlations from the ‘Cell-cycle process’ and ‘Mitotic cell cycle process’ themes. F) Distribution of correlation coefficients between mRNA and peptide abundances in small (Area<900um^2) cell lines. We note a positive skew and more positive mean than in the pooled distribution. G) Themes enriched in the set of genes exhibiting significant positive correlations between peptide and mRNA abundance in small cell lines. H) Example correlation between CCNB2 peptide and mRNA abundance from the ‘Mitotic cell cycle checkpoint’ theme. I) Distribution of correlation coefficients between mRNA and peptide abundances in big (Area>900um^2) cell lines. We note a negative skew and more negative mean than in the pooled distribution. J) Themes enriched in the set of genes exhibiting significant positive correlations between peptide and mRNA abundance in big cell lines. K) Example correlation between Rpl5 peptide and mRNA abundance from the ‘Translation’ theme. L) Expression of all detected RNA Pol1 (top left)/2 (right)/3 (bottom left) components across cell lines. Large lines tend to exhibit lower expression. M) Relationship between RNA Pol1/2/3 (left to right) peptide expression and the peptide (top) /mRNA (bottom) expression of identified RNA-peptide correlates in large cells in the ‘translation’ theme. We note that both RNA and peptide abundance correlate suggesting transcription regulation of peptide expression.

**Figure 6:**
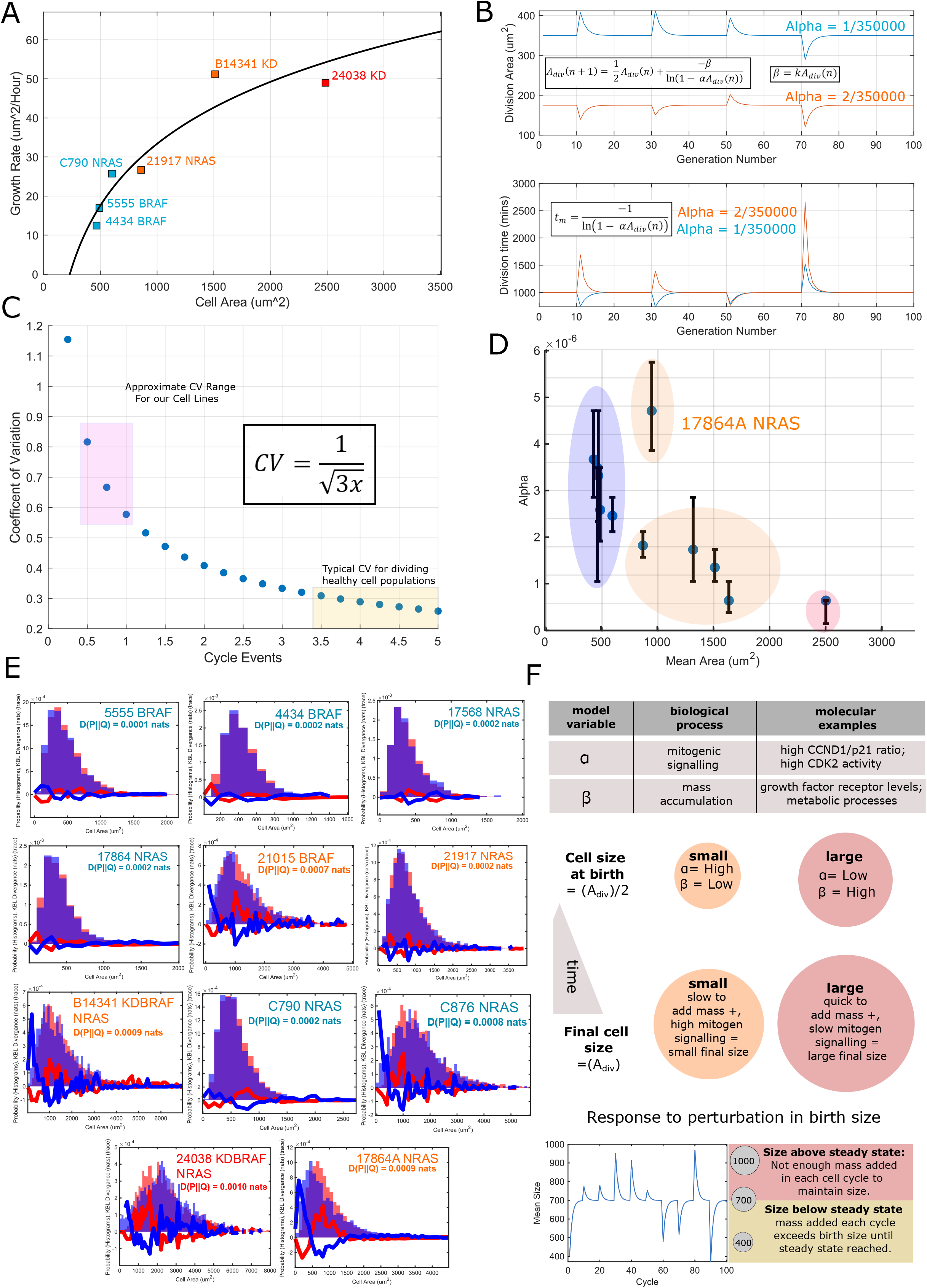
Theoretical modelling suggests decoupling of low mitogen and growth signalling drive proliferation of larger cells: A) The relationship between cell growth rate and cell size across six lines representing each genotype and size class. Growth rate increases with increasing cell size. B) A demonstration of the stability of the size distribution mean to perturbations to a cell division area and instability of the mean with respect to perturbations to alpha. The lower panel depicts the stability of proliferation rate with respect to both parameters C) The function describing how the coefficient of variation changes with increasing cycle complexity, the pink box marks the the CV’s observed in our cell lines and the yellow, those observed in other studies. D) The relationship between fitted alpha values and cell size, alpha broadly negatively correlates with the mean size of the cell line E) Model outputs demonstrating the best fits achieved. Orange histograms are model outputs and blue experimental data. The blue line shows D(experimental||measured) and the red line the reverse. F) Cartoon summary on the model relating parameters to biologically processes.

**Supplemental Figure 1:**
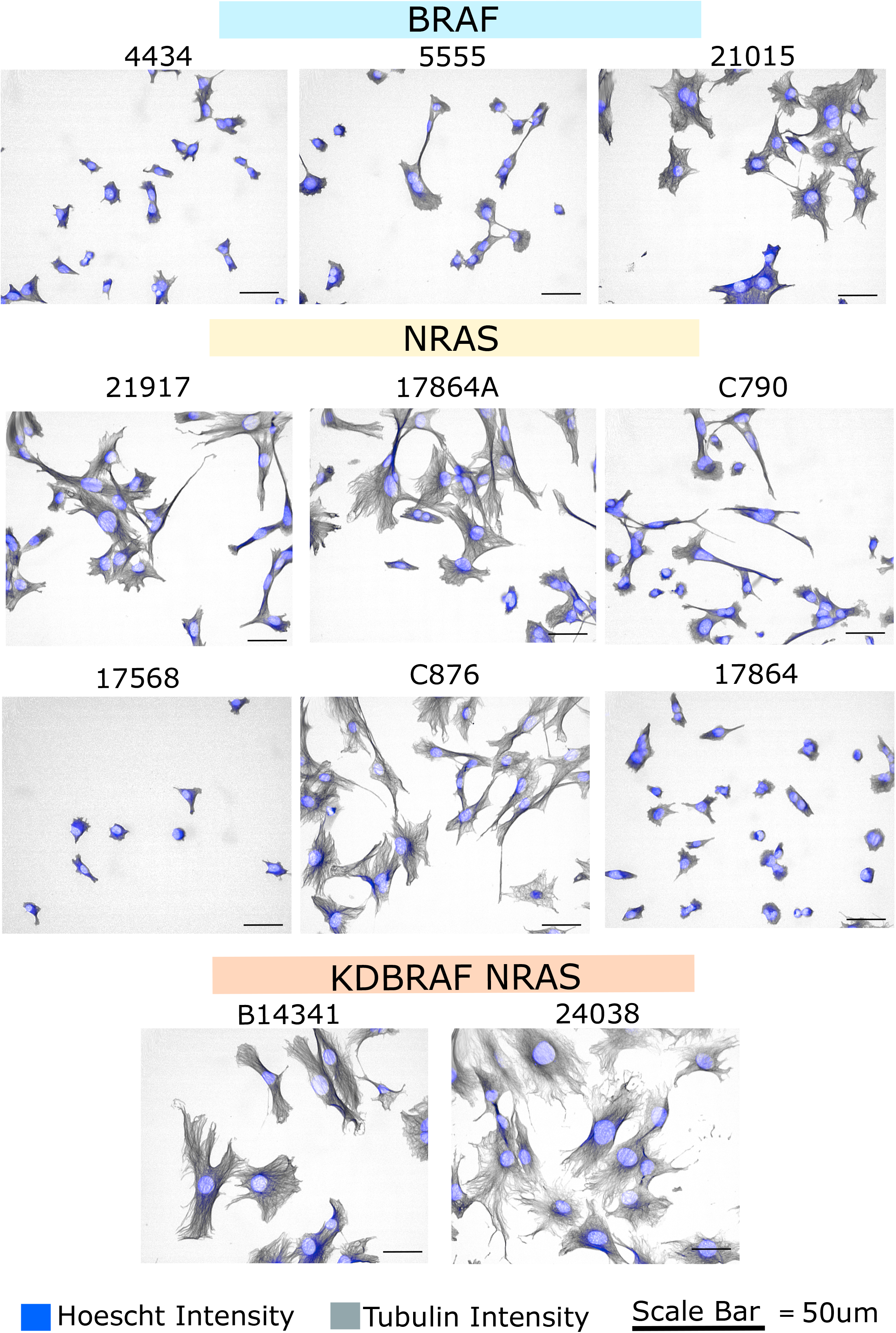
Images of the cell lines: A) Representative images from the 11 cell lines. In blue is the Hoechst intensity, and grey, the tubulin intensity. All images were taken at 20X magnification using an Opera Cell: Explorer-automated spinning disk confocal microscope. Images have been auto-adjusted to optimise contrast within the acapella environment (PerkinElmer).

**Supplemental Figure 2:**
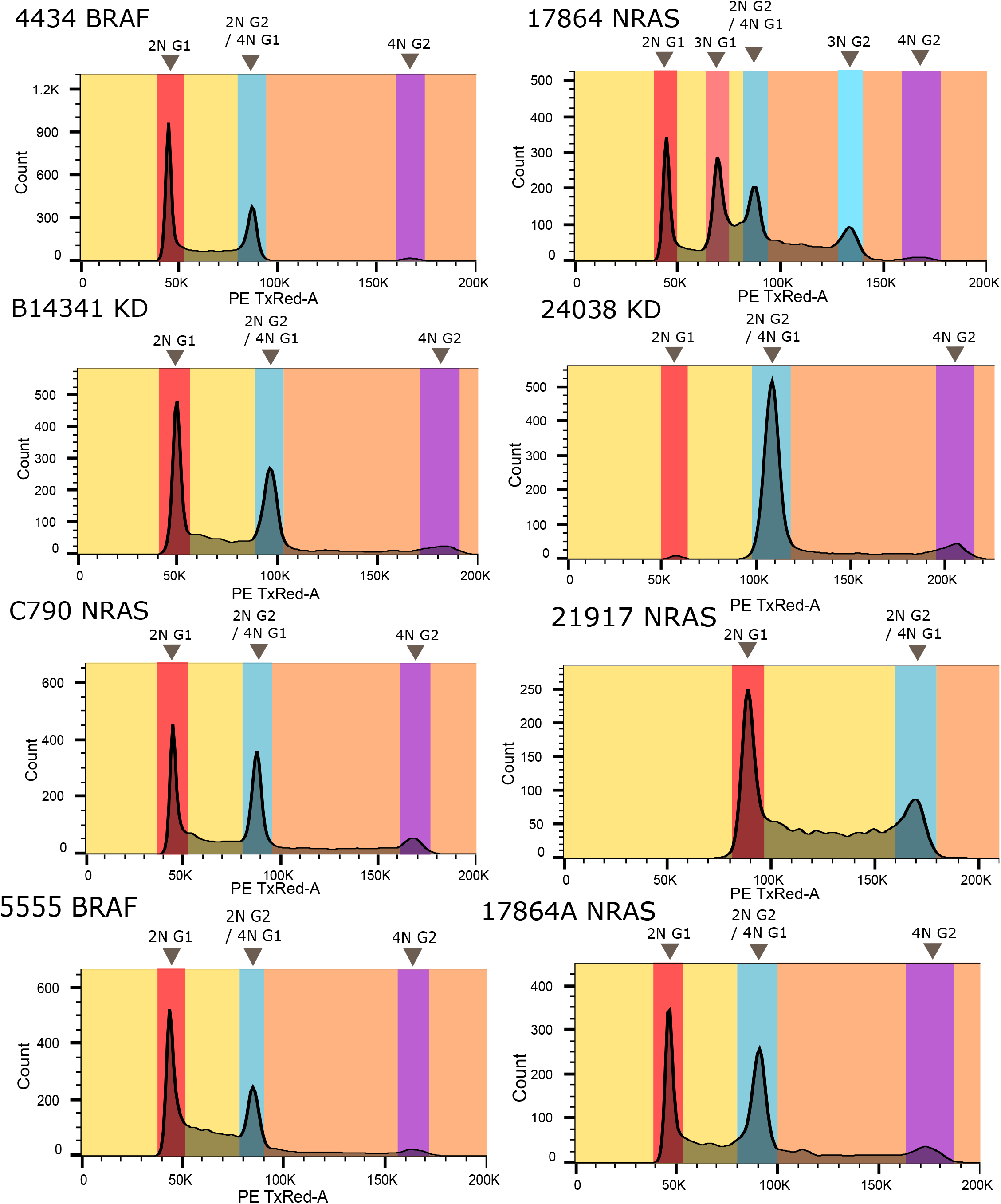
FACs Analysis: A) Quantification of cell DNA through FACs analysis. Many of the cell lines, both large and small, exhibit a small polyploid population. No clear relationship emerges between ploidy and cell size.

**Supplemental Figure 3:**
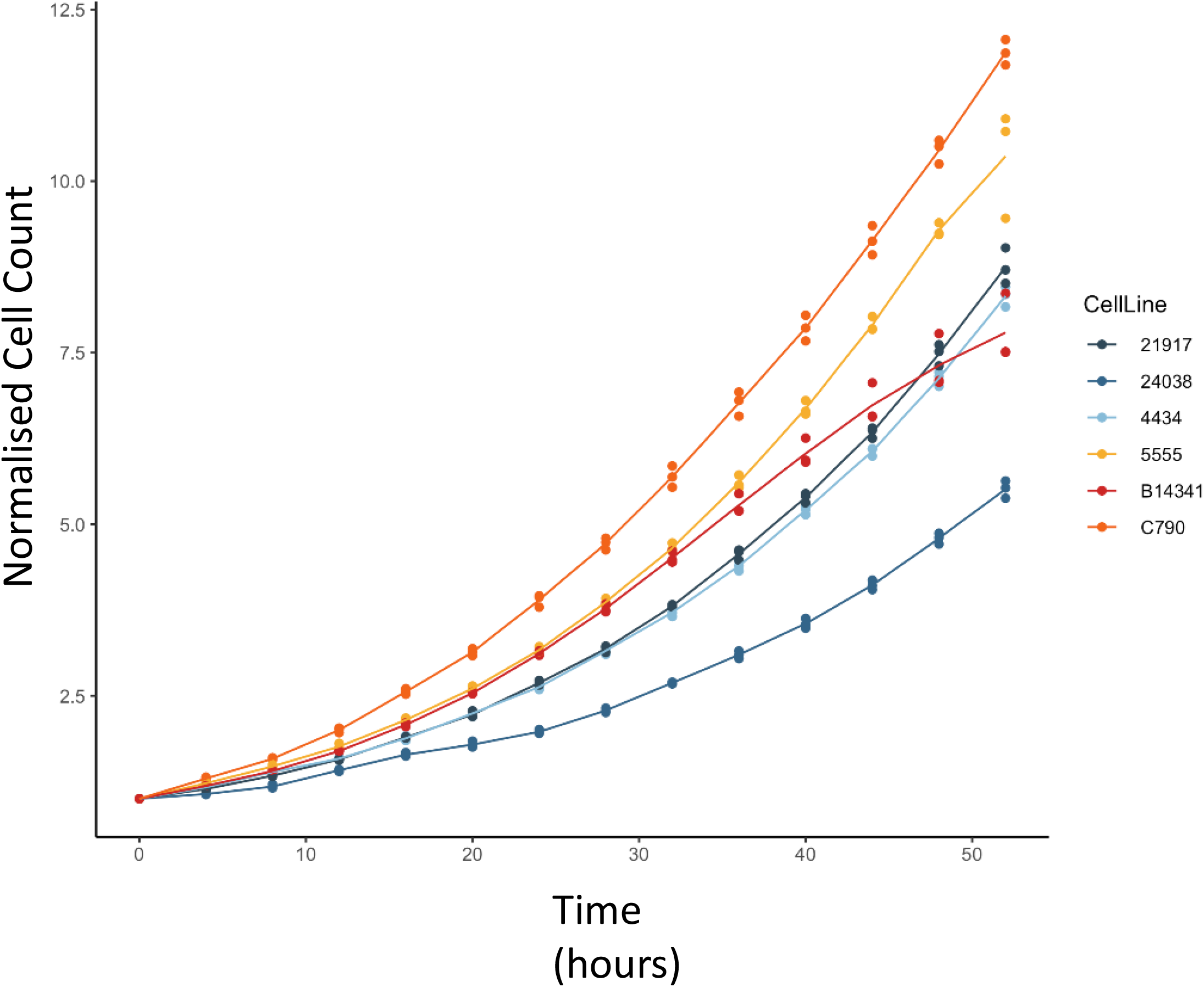
Cell growth data: Cell population growth curves for a subset of the investigated lines. Cell density is normalised relative to the starting confluence of the culture. Note that a large line, B14341, shows a comparable doubling time to smaller line, 5555.

**Supplemental figure 4:**
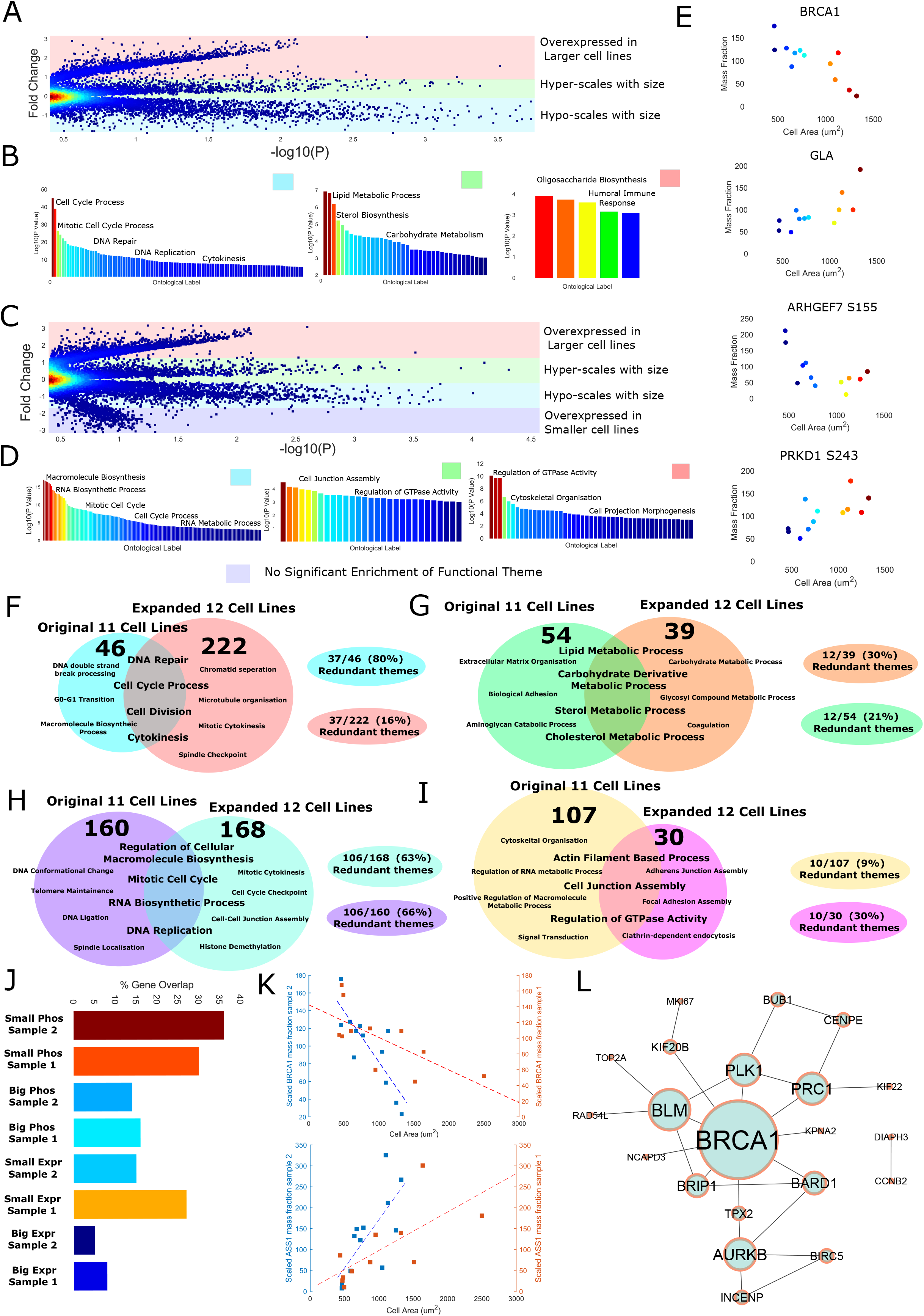
Validation of size controllers in an independent panel of melanoma cell lines: A) Volcano plot relating the ‘fold change’ across the small and large cell lines (defined as either side of the mean size) to the significance of the correlation between cell size. and peptide expression. B) Themes enriched in each region of ‘A’ as denoted by the colour of the box in the top right of each panel. From left to right; hypo-scales with size, hyper scales with size, over expressed in large cells. C) Volcano plot relating the ‘fold change’ across the small and large cell lines (defined as either side of the mean size) to the significance of the correlation between cell size and phosphopeptide expression. D) Themes enriched in each region of ‘C’ as denoted by the colour of the box in the top right of each panel. From left to right; hypo-scales with size, hyper scales with size, over expressed in large cells (bottom = overexpressed in small cells). E) Example hits from each analysis, the top half the peptide expression analysis, the bottom, the phosphopeptide expression analysis. F-I) Venn diagrams depicting the overlap of themes enriched across both sets of cell lines. Top left = peptide expression in small lines, top right = peptide expression in big lines, bottom left = phosphopeptide expression in small lines, bottom right, phosphopeptide expression in big lines. J) Percent overlap between analyses at the gene level. K) Example genes that are hits across both analyses, the top panel shows BRCA1 peptide expression and the bottom ASS1 peptide expression. L) A set of interacting peptides derived from the overlapping list of hit genes enriched in smaller cell lines centred on BRCA1.

**Supplemental Figure 5:**
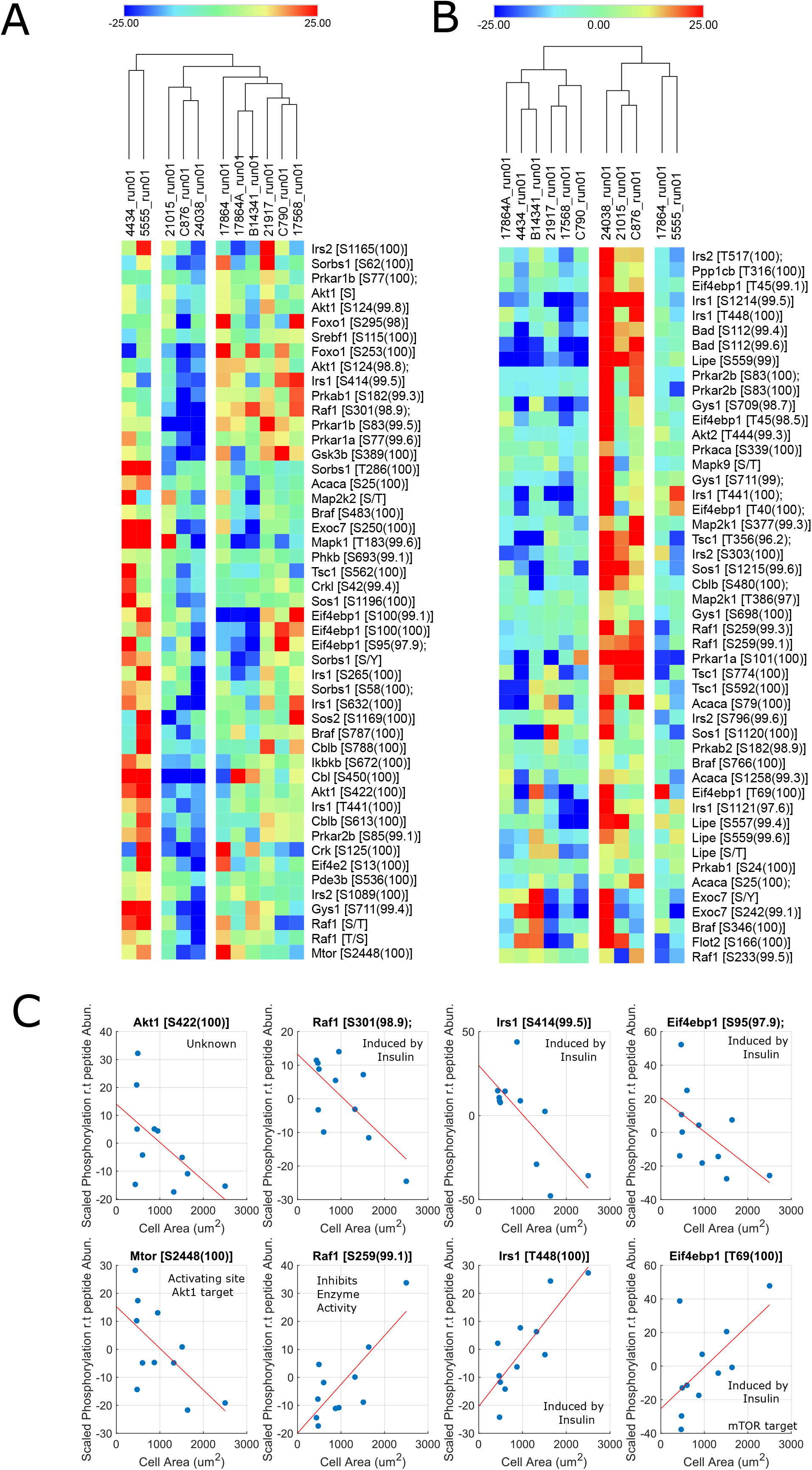
Growth signalling across cell lines: A) subset of phosphopeptides that negatively correlate with cell size pertaining to the ‘mTOR signalling’ KEGG pathway. B) As in ‘A’, but showing elements that positively correlate with cell size. C) Example correlation between mTOR signalling phosphorylations and cell size.

**Supplemental Figure 6:**
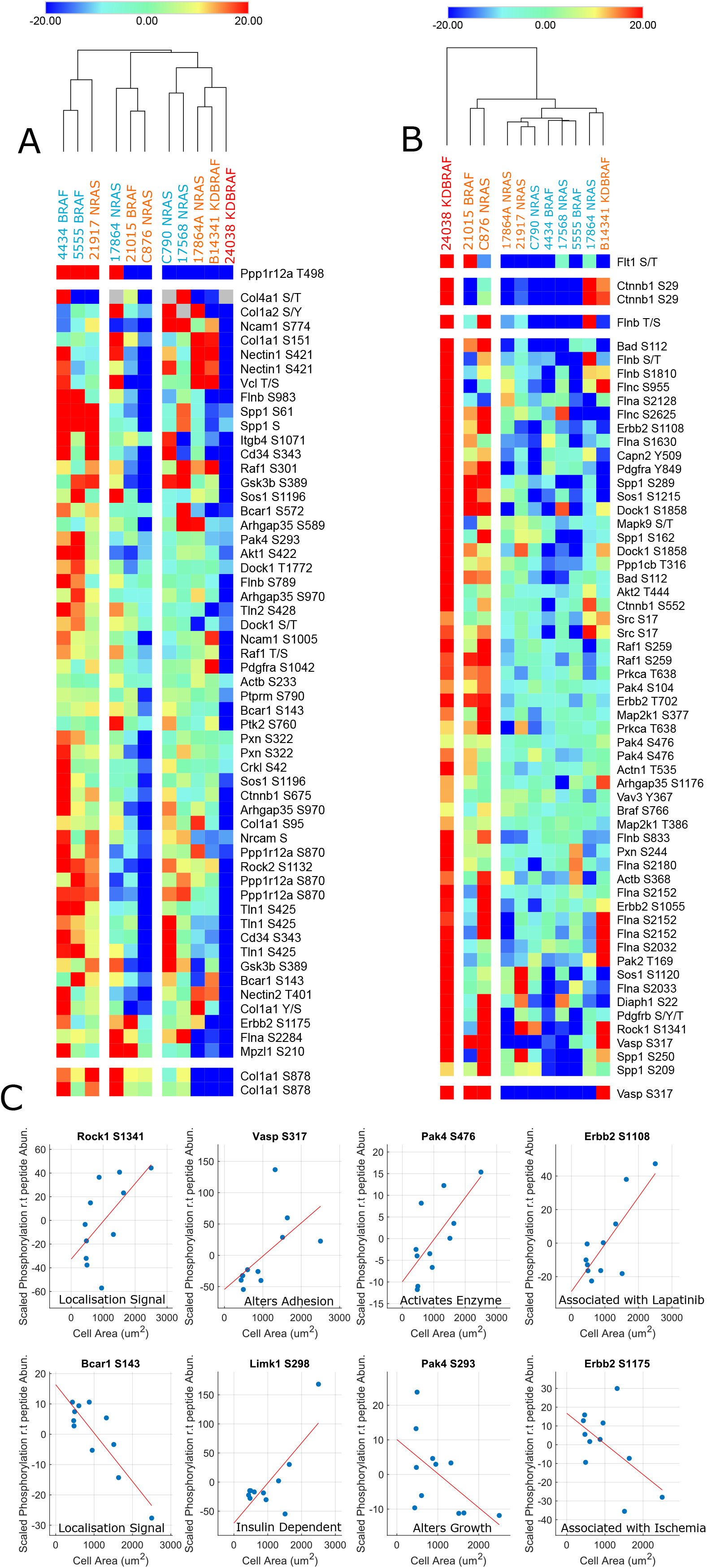
Cytoskeletal phosphorylation across cell lines: A) subset of phosphopeptides that negatively correlate with cell size pertaining to the ‘cytoskeleton’ and ‘adhesion’ KEGG pathways. B) As in ‘A’, but showing elements that positively correlate with cell size. C) Example correlation between cytoskeletal phosphorylations and cell size.

## Supplemental Information

### Analysis of gene and theme overlap of size-scaling factors between datasets

We investigated which ontological themes were enriched in both analyses finding that peptides pertaining to cell cycle, DNA repair, and division processes remained enriched in smaller cell lines (eg; ‘DNA repair’, ‘Cell cycle process’, ‘Cytokinesis’, 80% A1-A2, 16 % A2-A1 indicating 80% of themes enriched in the first analysis match the second and 16% detected in the second match the first) whilst lipid and carbohydrate metabolic peptides (eg; ‘Lipid metabolic process’, ‘Carbohydrate derivative metabolic process’, ‘Sterol metabolic process’, 30% A1-A2, 21% A2-A1) are consistently enriched in larger cell lines. Due to the lack of agreement, the enrichment of ECM components in larger cell lines detected in the prior analysis may reflect an upregulation or overexpression rather than a scaling relationship. Enacting the same analysis for the phosphorylation data, we note excellent agreement between analyses (63% A1 -A2, 60% A2-A1) for small cell lines, with both enriching for cell cycle and biosynthetic processes (eg; regulation of cellular biosynthetic process, mitotic cell cycle, DNA replication). Larger cell lines exhibited much weaker agreement (9% A1-A2, 30% A2-A1) but both analyses revealed enrichment of cytoskeletal and GTPase regulatory phosphorylations (eg; Regulation of GTPase activity, ‘Cell junction assembly’, ‘Actin filament based process’) (**SF4**).

Investigating the overlap of individual genes, we note a particularly strong overlap between analyses for phosphopeptides enriched in smaller cell lines (36% A1-A2, 30% A2-A1). Phosphopeptides enriched in larger cell lines show a more modest overlap (16% A1-A2, 14% A2-A1) like that observed in peptide expressions for smaller cell lines (15% A1-A2, 27% A2-A1). Peptide expressions in larger cell lines exhibit the weakest overlap (5% A1-A2, 8% A2-A1) (**SF3**). Screening for interactions between overlapping genes we observe a set of 21 physically interacting genes centred on BRCA1 enriched in smaller cell lines. As a ‘hit’ in two separate scaling analyses, these data indicate that the BRCA1 complex scales with cell size (**SF4**).

These data corroborate our previous analysis, strengthening the claim that G2/M and DNA repair processes define smaller melanoma cell lines, (with associated peptides sub-scaling with cell size), whilst cytoskeletal organisation and the rewiring of lipid metabolism define larger cell lines (peptides super-scaling with size). Interestingly we recover a large, BRCA1 complex in both analyses, implicating the complex in size-dependent phenomena.

### Derivation of the proliferation time and size gain distributions

We are interested in the waiting time distribution before the first successful event. The probability to fail a division is:

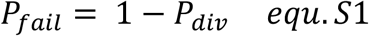

For a cell to have not divided by a given time point, it must have failed to divide at every prior time point. The probability of successive failures occurring at a given time is equal to:

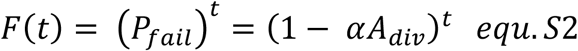

Where ‘t’ is time since the last division. The probability of having divided by a given ‘t’ is the probability that the cell has not failed at every prior step:

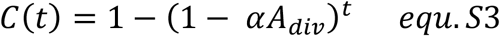

The probability distribution follows as:

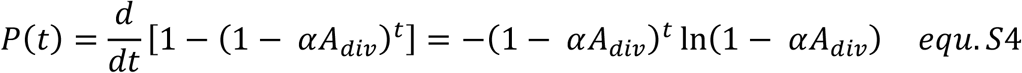

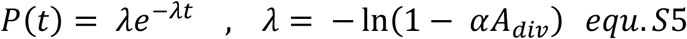

We may extract the expected gained mass by scaling the time by ln(2)/(dt/dA). The ln(2) factor accounts for a division event having happened any time in the interval 0-t.

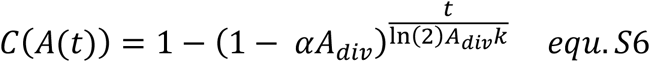

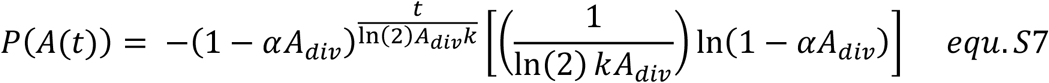

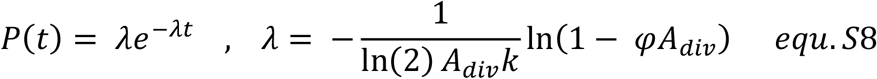

With a mean of known form given as 1/ *λ* :

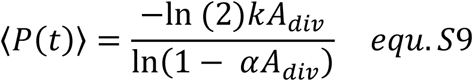

We can see that this result constitutes an adder –type system when expressing the expected area gain as a Laurent series about *α* = 0 (fitted values never exceed 1×10^-5) (F6B/C, table 2):

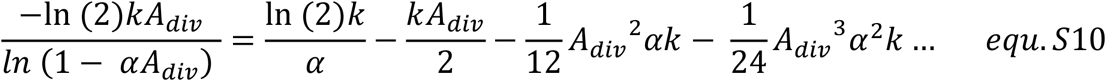

The mean area gain is approximately constant, as the first term dominates the expression by virtue of alpha ≈ 0. Thus, a constant average mass is added each cycle, despite the area gain distribution itself being dependent on division size.

### Deriving the moments of the cell size distribution

Starting with an initial size distribution, F(A), and size gain distribution, G(A), we may define the expected size distribution up to the first division, H(A) as:

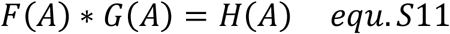

On division, the value of cell size is considered to halve. Thus, the birth size distribution is given as:

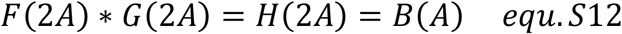

Where the inclusion of 2A has mapped the probability of A to half its value, thereby simulating a division event. This is then convolved with G(A) again for the next division cycle, and so on:

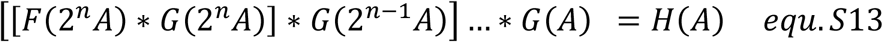

Where n denotes the number of divisions. Note that as n increases, the influence of the initial size distribution on the total convolution decreases as *F*(2^*n*^*A*) has non-zeros values only at extremely low sizes as n increases. Indeed, we can approximate the above as:

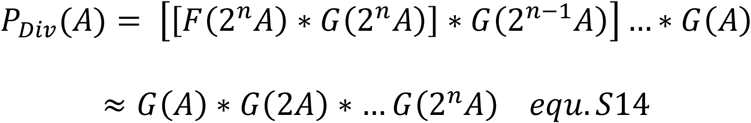

G(A) has been shown to be an exponential distribution. Convolution of n exponential functions with different scale parameters results in a hypo-exponential function with mean equal to the sum of the means of all participating distributions:

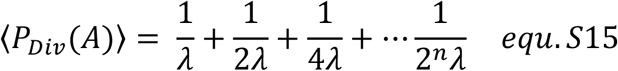

The sum can be written as:

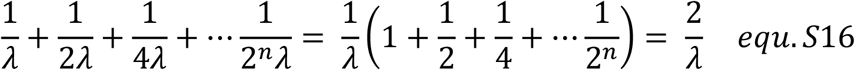

Indicating that the distribution tends toward a constant mean. The corresponding variance is similarly given as:

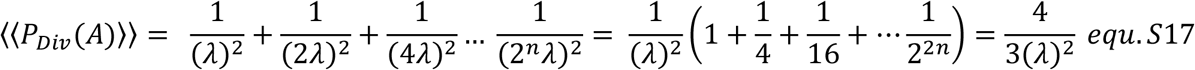

Yielding a constant coefficient of variation:

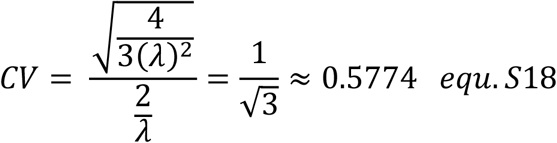

These results may be trivially adjusted to account for ‘x’ identical events governing division. Indeed, G(A) is merely transformed from a constant exponential distribution to a constant Erlang distribution of shape factor ‘x’ and rate parameter 1/x k/a. This stems from G(A) being generated from the convolution of ‘x’ exponentially distributed gain variables corresponding to the area gain in each cycle stage each with mean 1/(x) k/a. As is the case for the hypoexponential, Erlang distributions have means and variance equal to the sum of those of the participating distributions allowing us to easily modify equ.S16/17:

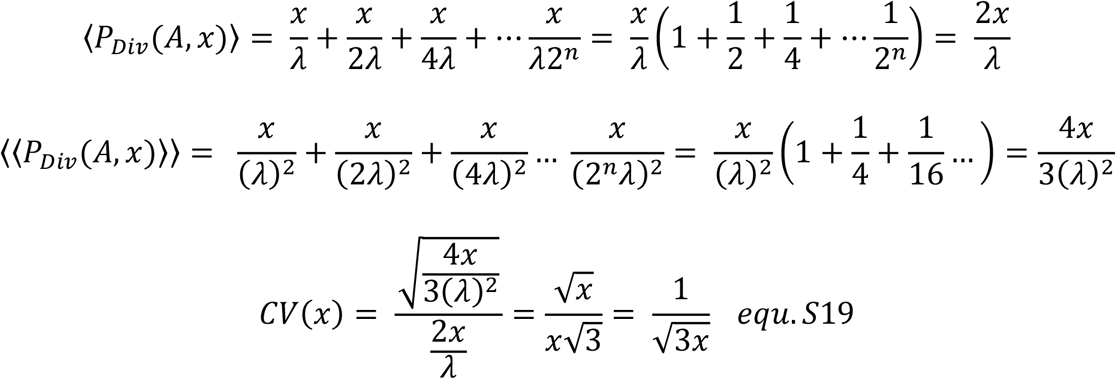

Equ.S19 tells us that from the coefficient of variation, we may estimate the number of stages needed to effectively model the cell size distributions. This relationship is similar to that obtained recently (Nieto et al., 2020) where (CV)^2 was found to be proportional to one over the number of modelled cell cycle stages. Importantly, given a single value of the ′*α*^′^ or ‘k’ parameters, this is entirely independent of ′*α*^′^ or ‘k’ facilitating simple calculation of the required ‘x’:

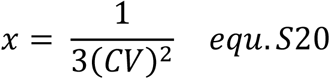

## References

Amodeo, A. A., Jukam, D., Straight, A. F., & Skotheim, J. M. (2015). Histone titration against the genome sets the DNA-to-cytoplasm threshold for the Xenopus midblastula transition. Proceedings of the National Academy of Sciences of the United States of America, 112(10). https://doi.org/10.1073/pnas.1413990112

Amodeo, A. A., & Skotheim, J. M. (2016). Cell-size control. Cold Spring Harbor Perspectives in Biology, 8(4). https://doi.org/10.1101/cshperspect.a019083

Andrew, A. M. (2004). Information Theory, Inference, and Learning Algorithms. In Kybernetes (Vol. 33, Issue 7). https://doi.org/10.1108/03684920410534506

Ashburner, M., Ball, C. A., Blake, J. A., Botstein, D., Butler, H., Cherry, J. M., Davis, A. P., Dolinski, K., Dwight, S. S., Eppig, J. T., Harris, M. A., Hill, D. P., Issel-Tarver, L., Kasarskis, A., Lewis, S., Matese, J. C., Richardson, J. E., Ringwald, M., Rubin, G. M., & Sherlock, G. (2000). Gene ontology: Tool for the unification of biology. In Nature Genetics (Vol. 25, Issue 1). https://doi.org/10.1038/75556

Bakal, C., Aach, J., Church, G., & Perrimon, N. (2007). Quantitative morphological signatures define local signaling networks regulating cell morphology. Science, 316(5832). https://doi.org/10.1126/science.1140324

Baryshnikova, A. (2016). Systematic Functional Annotation and Visualization of Biological Networks. Cell Systems, 2(6). https://doi.org/10.1016/j.cels.2016.04.014

Bernal-Mizrachi, E., Wen, W., Stahlhut, S., Welling, C. M., & Permutt, M. A. (2001). Islet β cell expression of constitutively active Akt1/PKBα induces striking hypertrophy, hyperplasia, and hyperinsulinemia. Journal of Clinical Investigation, 108(11). https://doi.org/10.1172/jci13785

Bhargava, A., Anant, M., Mack, H., Iorns, E., Gunn, W., Tan, F., Lomax, J., Williams, S. R., Perfito, N., & Errington, T. (2016). Registered report: Kinase-dead BRAF and oncogenic RAS cooperate to drive tumor progression through CRAF. ELife, 5(FEBRUARY 2016). https://doi.org/10.7554/eLife.11999

Burd, C. E., Liu, W., Huynh, M. v., Waqas, M. A., Gillahan, J. E., Clark, K. S., Fu, K., Martin, B. L., Jeck, W. R., Souroullas, G. P., Darr, D. B., Zedek, D. C., Miley, M. J., Baguley, B. C., Campbell, S. L., & Sharpless, N. E. (2014). Mutation-specific RAS oncogenicity explains NRAS codon 61 selection in melanoma. Cancer Discovery, 4(12), 1418–1429. https://doi.org/10.1158/2159-8290.CD-14-0729

Campos, M., Surovtsev, I. v., Kato, S., Paintdakhi, A., Beltran, B., Ebmeier, S. E., & Jacobs-Wagner, C. (2014). A constant size extension drives bacterial cell size homeostasis. Cell, 159(6). https://doi.org/10.1016/j.cell.2014.11.022

Cantwell-Dorris, E. R., O’Leary, J. J., & Sheils, O. M. (2011). BRAFV600E: Implications for carcinogenesis and molecular therapy. In Molecular Cancer Therapeutics (Vol. 10, Issue 3). https://doi.org/10.1158/1535-7163.MCT-10-0799

Carbon, S., Douglass, E., Good, B. M., Unni, D. R., Harris, N. L., Mungall, C. J., Basu, S., Chisholm, R. L., Dodson, R. J., Hartline, E., Fey, P., Thomas, P. D., Albou, L. P., Ebert, D., Kesling, M. J., Mi, H., Muruganujan, A., Huang, X., Mushayahama, T., … Elser, J. (2021). The Gene Ontology resource: Enriching a GOld mine. Nucleic Acids Research, 49(D1). https://doi.org/10.1093/nar/gkaa1113

Caspersson, T., Foley, G. E., Killander, D., & Lomakka, G. (1963). Cytochemical differences between mammalian cell lines of normal and neoplastic origins. Experimental Cell Research, 32(3). https://doi.org/10.1016/0014-4827(63)90193-9

Chao, H. X., Fakhreddin, R. I., Shimerov, H. K., Kedziora, K. M., Kumar, R. J., Perez, J., Limas, J. C., Grant, G. D., Cook, J. G., Gupta, G. P., & Purvis, J. E. (2019). Evidence that the human cell cycle is a series of uncoupled, memoryless phases. Molecular Systems Biology, 15(3). https://doi.org/10.15252/msb.20188604

Chiang, G. G., & Abraham, R. T. (2005). Phosphorylation of mammalian target of rapamycin (mTOR) at Ser-2448 is mediated by p70S6 kinase. Journal of Biological Chemistry, 280(27). https://doi.org/10.1074/jbc.M501707200

Chinnam, M., & Goodrich, D. W. (2011). RB1, Development, and Cancer. In Current Topics in Developmental Biology (Vol. 94, Issue C). https://doi.org/10.1016/B978-0-12-380916-2.00005-X

Costanzo, M., Nishikawa, J. L., Tang, X., Millman, J. S., Schub, O., Breitkreuz, K., Dewar, D., Rupes, I., Andrews, B., & Tyers, M. (2004). CDK activity antagonizes Whi5, an inhibitor of G1/S transcription in yeast. Cell, 117(7). https://doi.org/10.1016/j.cell.2004.05.024

Davies, H., Bignell, G. R., Cox, C., Stephens, P., Edkins, S., Clegg, S., Teague, J., Woffendin, H., Garnett, M. J., Bottomley, W., Davis, N., Dicks, E., Ewing, R., Floyd, Y., Gray, K., Hall, S., Hawes, R., Hughes, J., Kosmidou, V., … Futreal, P. A. (2002). Mutations of the BRAF gene in human cancer. Nature, 417(6892). https://doi.org/10.1038/nature00766

Dhomen, N., Reis-Filho, J. S., da Rocha Dias, S., Hayward, R., Savage, K., Delmas, V., Larue, L., Pritchard, C., & Marais, R. (2009). Oncogenic Braf Induces Melanocyte Senescence and Melanoma in Mice. Cancer Cell, 15(4). https://doi.org/10.1016/j.ccr.2009.02.022

Dunham, I., Kundaje, A., Aldred, S. F., Collins, P. J., Davis, C. A., Doyle, F., Epstein, C. B., Frietze, S., Harrow, J., Kaul, R., Khatun, J., Lajoie, B. R., Landt, S. G., Lee, B. K., Pauli, F., Rosenbloom, K. R., Sabo, P., Safi, A., Sanyal, A., … Lochovsky, L. (2012). An integrated encyclopedia of DNA elements in the human genome. Nature, 489(7414). https://doi.org/10.1038/nature11247

Eden, E., Navon, R., Steinfeld, I., Lipson, D., & Yakhini, Z. (2009). GOrilla: A tool for discovery and visualization of enriched GO terms in ranked gene lists. BMC Bioinformatics, 10. https://doi.org/10.1186/1471-2105-10-48

Facchetti, G., Knapp, B., Chang, F., & Howard, M. (2019). Reassessment of the Basis of Cell Size Control Based on Analysis of Cell-to-Cell Variability. Biophysical Journal, 117(9). https://doi.org/10.1016/j.bpj.2019.09.031

Facchetti, G., Knapp, B., Flor-Parra, I., Chang, F., & Howard, M. (2019). Reprogramming Cdr2-Dependent Geometry-Based Cell Size Control in Fission Yeast. Current Biology, 29(2). https://doi.org/10.1016/j.cub.2018.12.017

Fantes, P., & Nurse, P. (1977). Control of cell size at division in fission yeast by a growth-modulated size control over nuclear division. Experimental Cell Research, 107(2). https://doi.org/10.1016/0014-4827(77)90359-7

Fingar, D. C., Richardson, C. J., Tee, A. R., Cheatham, L., Tsou, C., & Blenis, J. (2004). mTOR Controls Cell Cycle Progression through Its Cell Growth Effectors S6K1 and 4E-BP1/Eukaryotic Translation Initiation Factor 4E. Molecular and Cellular Biology, 24(1). https://doi.org/10.1128/mcb.24.1.200-216.2004

Franceschini, A., Szklarczyk, D., Frankild, S., Kuhn, M., Simonovic, M., Roth, A., Lin, J., Minguez, P., Bork, P., von Mering, C., & Jensen, L. J. (2013). STRING v9.1: Protein-protein interaction networks, with increased coverage and integration. Nucleic Acids Research, 41(D1). https://doi.org/10.1093/nar/gks1094

Gennaro, V. J., Stanek, T. J., Peck, A. R., Sun, Y., Wang, F., Qie, S., Knudsen, K. E., Rui, H., Butt, T., Diehl, J. A., & McMahon, S. B. (2018). Control of CCND1 ubiquitylation by the catalytic SAGA subunit USP22 is essential for cell cycle progression through G1 in cancer cells. Proceedings of the National Academy of Sciences of the United States of America, 115(40). https://doi.org/10.1073/pnas.1807704115

Ginzberg, M. B., Kafri, R., & Kirschner, M. (2015). On being the right (cell) size. In Science (Vol. 348, Issue 6236). https://doi.org/10.1126/science.1245075

Gonzalez, N. P., Tao, J., Rochman, N. D., Vig, D., Chiu, E., Wirtz, D., & Sun, S. X. (2018). Cell tension and mechanical regulation of cell volume. Molecular Biology of the Cell, 29(21). https://doi.org/10.1091/mbc.E18-04-0213

Gry, M., Rimini, R., Strömberg, S., Asplund, A., Pontén, F., Uhlén, M., & Nilsson, P. (2009). Correlations between RNA and protein expression profiles in 23 human cell lines. BMC Genomics, 10. https://doi.org/10.1186/1471-2164-10-365

Guo, M., Pegoraro, A. F., Mao, A., Zhou, E. H., Arany, P. R., Han, Y., Burnette, D. T., Jensen, M. H., Kasza, K. E., Moore, J. R., Mackintosh, F. C., Fredberg, J. J., Mooney, D. J., Lippincott-Schwartz, J., & Weitz, D. A. (2017). Cell volume change through water efflux impacts cell stiffness and stem cell fate. Proceedings of the National Academy of Sciences of the United States of America, 114(41). https://doi.org/10.1073/pnas.1705179114

Harris, L. K., & Theriot, J. A. (2016). Relative rates of surface and volume synthesis set bacterial cell size. Cell, 165(6). https://doi.org/10.1016/j.cell.2016.05.045

Heldt, F. S., Lunstone, R., Tyson, J. J., & Novák, B. (2018). Dilution and titration of cell-cycle regulators may control cell size in budding yeast. PLoS Computational Biology, 14(10). https://doi.org/10.1371/journal.pcbi.1006548

Hornbeck, P. v., Zhang, B., Murray, B., Kornhauser, J. M., Latham, V., & Skrzypek, E. (2015). PhosphoSitePlus, 2014: Mutations, PTMs and recalibrations. Nucleic Acids Research, 43(D1). https://doi.org/10.1093/nar/gku1267

Joseph, E. W., Pratilas, C. A., Poulikakos, P. I., Tadi, M., Wang, W., Taylor, B. S., Halilovic, E., Persaud, Y., Xing, F., Viale, A., Tsai, J., Chapman, P. B., Bollag, G., Solit, D. B., & Rosen, N. (2010). The RAF inhibitor PLX4032 inhibits ERK signaling and tumor cell proliferation in a V600E BRAF-selective manner. Proceedings of the National Academy of Sciences of the United States of America, 107(33). https://doi.org/10.1073/pnas.1008990107

Kim, M. H., Kim, J., Hong, H., Lee, S., Lee, J., Jung, E., & Kim, J. (2016). Actin remodeling confers BRAF inhibitor resistance to melanoma cells through YAP / TAZ activation. The EMBO Journal, 35(5). https://doi.org/10.15252/embj.201592081

Lanz, M. C., Zatulovskiy, E., Swaffer, M. P., Zhang, L., Ilerten, I., Zhang, S., You, D. S., Marinov, G., McAlpine, P., Elias, J. E., & Skotheim, J. M. (2021). Increasing cell size remodels the proteome and promotes senescence. BioRxiv.

Lin, L., & Bivona, T. G. (2016). The Hippo effector YAP regulates the response of cancer cells to MAPK pathway inhibitors. Molecular and Cellular Oncology, 3(1). https://doi.org/10.1080/23723556.2015.1021441

Lundgren, K., Walworth, N., Booher, R., Dembski, M., Kirschner, M., & Beach, D. (1991). mik1 and wee1 cooperate in the inhibitory tyrosine phosphorylation of cdc2. Cell, 64(6). https://doi.org/10.1016/0092-8674(91)90266-2

Malin Pedersen, Amaya Viros, Martin Cook, & Richard Marais. (2014). (G12D) NRAS and kinase-dead BRAF cooperate to drive naevogenesis and melanomagenesis. Pigment Cell Melanoma Res., 27(6), 1162–1166.

Mi, H., Muruganujan, A., Ebert, D., Huang, X., & Thomas, P. D. (2019). PANTHER version 14: More genomes, a new PANTHER GO-slim and improvements in enrichment analysis tools. Nucleic Acids Research, 47(D1). https://doi.org/10.1093/nar/gky1038

Miettinen, T. P., Ly, K. S., Lam, A., & Manalis, S. R. (2021). Single-cell monitoring of dry mass and dry density reveals exocytosis of cellular dry contents in mitosis. BioRxiv.

Min, M., Rong, Y., Tian, C., & Spencer, S. L. (2020). Temporal integration of mitogen history in mother cells controls proliferation of daughter cells. Science, 368(6496). https://doi.org/10.1126/science.aay8241

Monds, R. D., Lee, T. K., Colavin, A., Ursell, T., Quan, S., Cooper, T. F., & Huang, K. C. (2014). Systematic Perturbation of Cytoskeletal Function Reveals a Linear Scaling Relationship between Cell Geometry and Fitness. Cell Reports, 9(4). https://doi.org/10.1016/j.celrep.2014.10.040

Neurohr, G. E., Terry, R. L., Lengefeld, J., Bonney, M., Brittingham, G. P., Moretto, F., Miettinen, T. P., Vaites, L. P., Soares, L. M., Paulo, J. A., Harper, J. W., Buratowski, S., Manalis, S., van Werven, F. J., Holt, L. J., & Amon, A. (2019). Excessive Cell Growth Causes Cytoplasm Dilution And Contributes to Senescence. Cell, 176(5). https://doi.org/10.1016/j.cell.2019.01.018

Nevins, J. R. (2001). The Rb/E2F pathway and cancer. In Human Molecular Genetics (Vol. 10, Issue 7). https://doi.org/10.1093/hmg/10.7.699

Nieto, C., Arias-Castro, J., Sánchez, C., Vargas-Garciá, C., & Pedraza, J. M. (2020). Unification of cell division control strategies through continuous rate models. Physical Review E, 101(2). https://doi.org/10.1103/PhysRevE.101.022401

Palenik, B., Grimwood, J., Aerts, A., Rouzé, P., Salamov, A., Putnam, N., Dupont, C., Jorgensen, R., Derelle, E., Rombauts, S., Zhou, K., Otillar, R., Merchant, S. S., Podell, S., Gaasterland, T., Napoli, C., Gendler, K., Manuell, A., Tai, V., … Grigoriev, I. v. (2007). The tiny eukaryote Ostreococcus provides genomic insights into the paradox of plankton speciation. Proceedings of the National Academy of Sciences of the United States of America, 104(18). https://doi.org/10.1073/pnas.0611046104

Pedersen, M., Küsters-Vandevelde, H. V. N., Viros, A., Groenen, P. J. T. A., Sanchez-Laorden, B., Gilhuis, J. H., van Engen-van Grunsven, I. A., Renier, W., Schieving, J., Niculescu-Duvaz, I., Springer, C. J., Küsters, B., Wesseling, P., Blokx, W. A. M., & Marais, R. (2013). Primary melanoma of the CNS in children is driven by congenital expression of oncogenic NRAS in melanocytes. Cancer Discovery, 3(4). https://doi.org/10.1158/2159-8290.CD-12-0464

Pedersen, M., Viros, A., & Marais, R. (2014). Abstract B11: Mouse models of melanoma driven by oncogenic NRAS. https://doi.org/10.1158/1557-3125.rasonc14-b11

Poulikakos, P. I., Zhang, C., Bollag, G., Shokat, K. M., & Rosen, N. (2010). RAF inhibitors transactivate RAF dimers and ERK signalling in cells with wild-type BRAF. Nature, 464(7287). https://doi.org/10.1038/nature08902

Reifenberger, J., Knobbe, C. B., Sterzinger, A. A., Blaschke, B., Schulte, K. W., Ruzicka, T., & Reifenberger, G. (2004). Frequent alterations of Ras signaling pathway genes in sporadic malignant melanomas. International Journal of Cancer, 109(3). https://doi.org/10.1002/ijc.11722

Russell, P., & Nurse, P. (1987). Negative regulation of mitosis by wee1+, a gene encoding a protein kinase homolog. Cell, 49(4). https://doi.org/10.1016/0092-8674(87)90458-2

Ruvinsky, I., Sharon, N., Lerer, T., Cohen, H., Stolovich-Rain, M., Nir, T., Dor, Y., Zisman, P., & Meyuhas, O. (2005). Ribosomal protein S6 phosphorylation is a determinant of cell size and glucose homeostasis. Genes and Development, 19(18). https://doi.org/10.1101/gad.351605

Schmoller, K. M., Turner, J. J., Kõivomägi, M., & Skotheim, J. M. (2015). Dilution of the cell cycle inhibitor Whi5 controls budding-yeast cell size. Nature, 526(7572). https://doi.org/10.1038/nature14908

Scotchman, E., Kume, K., Navarro, F. J., & Nurse, P. (2021). Identification of mutants with increased variation in cell size at onset of mitosis in fission yeast. Journal of Cell Science, 134(3). https://doi.org/10.1242/jcs.251769

Serbanescu, D., Ojkic, N., & Banerjee, S. (2020). Nutrient-Dependent Trade-Offs between Ribosomes and Division Protein Synthesis Control Bacterial Cell Size and Growth. Cell Reports, 32(12). https://doi.org/10.1016/j.celrep.2020.108183

Shannon, P., Markiel, A., Ozier, O., Baliga, N. S., Wang, J. T., Ramage, D., Amin, N., Schwikowski, B., & Ideker, T. (2003). Cytoscape: A software Environment for integrated models of biomolecular interaction networks. Genome Research, 13(11). https://doi.org/10.1101/gr.1239303

Stallaert, W., Kedziora, K. M., Taylor, C. D., Zikry, T. M., Sobon, H. K., Taylor, S. R., Young, C. L., Limas, J. C., Cook, J. G., & Purvis, J. E. (2021). The structure of the human cell cycle. BioRxiv, 2021.02.11.430845. http://biorxiv.org/content/early/2021/02/11/2021.02.11.430845.abstract

Swaffer, M. P., Kim, J., Chandler-Brown, D., Langhinrichs, M., Marinov, G. K., Greenleaf, W. J., Kundaje, A., Schmoller, K. M., & Skotheim, J. M. (2021). Transcriptional and chromatin-based partitioning mechanisms uncouple protein scaling from cell size. Molecular Cell, 81(23). https://doi.org/10.1016/j.molcel.2021.10.007

Varsano, G., Wang, Y., & Wu, M. (2017). Probing Mammalian Cell Size Homeostasis by Channel-Assisted Cell Reshaping. Cell Reports, 20(2). https://doi.org/10.1016/j.celrep.2017.06.057

Venkova, L., Vishen, A. S., Lembo, S., Srivastava, N., Duchamp, B., Ruppel, A., Vassilopoulos, S., Deslys, A., Arcos, J. M. G., Diz-Muñoz, A., Balland, M., Joanny, J.-F., Cuvelier, D., Sens, P., & Piel, M. (2021). A mechano-osmotic feedback couples cell volume to the rate of cell deformation. BioRxiv. https://doi.org/10.1101/2021.06.08.447538

Vízkeleti, L., Ecsedi, S., Rákosy, Z., Orosz, A., Lázár, V., Emri, G., Koroknai, V., Kiss, T., Ádány, R., & Balázs, M. (2012). The role of CCND1 alterations during the progression of cutaneous malignant melanoma. Tumour Biology : The Journal of the International Society for Oncodevelopmental Biology and Medicine, 33(6). https://doi.org/10.1007/s13277-012-0480-6

Watson, G. (1997). Cells, Tissues and Disease. Principles of General Pathology. Pathology, 29(3). https://doi.org/10.1080/00313029700169265

Xie, K., Yang, Y., & Jiang, H. (2018). Controlling Cellular Volume via Mechanical and Physical Properties of Substrate. Biophysical Journal, 114(3). https://doi.org/10.1016/j.bpj.2017.11.3785

Yahya, G., Menges, P., Ngandiri, D. A., Schulz, D., Wallek, A., Kulak, N., Mann, M., Cramer, P., Savage, V., Raeschle, M., Storchova, Z., Yahya, G., Menges, P., Ngandiri, D. A., Schulz, D., Wallek, A., Kulak, N., Mann, M., Cramer, P., … Storchova, Z. (2021). Scaling of cellular proteome with ploidy. BioRxiv.

Zatulovskiy, E., Zhang, S., Berenson, D. F., Topacio, B. R., & Skotheim, J. M. (2020). Cell growth dilutes the cell cycle inhibitor Rb to trigger cell division. Science, 369(6502). https://doi.org/10.1126/science.aaz6213

